# C3 and CD47 mediate sensory-motor circuit refinement during spinal cord development

**DOI:** 10.1101/2023.11.08.566259

**Authors:** D Florez-Paz, G.Z. Mentis

## Abstract

Overground movement in mammals requires the assembly and refinement of sensory-motor circuits. Within the spinal cord, sensory neurons, interneurons and motor neurons form intricate neuronal circuits to ensure proper motor control. In the developing brain, supernumerary synapses are initially formed and subsequently pruned making way for the emergence of mature circuits. However, whether this occurs within spinal sensory-motor circuits, it has not been firmly established. Moreover, it is also unknown if a combination of distinct molecules are required to refine spinal cord neuronal circuits. Here, we demonstrate the presence of supernumerary synapses which form inappropriate contacts, resulting in miswired immature spinal neuronal circuits. We determined that inappropriate synapses are of proprioceptive sensory origin and are functional, leading to impaired motor behavior. Using mouse genetics, viral-mediated neuronal map strategies, electrophysiology, and behavioral assessments, we demonstrate that two molecularly distinct mechanisms are responsible for the refinement of spinal circuits. First, we identify C3 as a major contributing factor through classical complement activation. Second, a CD47-dependent mechanism, operating in parallel to classical complement, causing elimination of inappropriate synapses. This finding underlies an unexpected function for CD47 within the spinal cord, in striking contrast to its function in the brain. Our study demonstrates that during early development, the natural course of elimination of inappropriately-generated synapses utilizes a dual fail-safe system to ensure the emergence of normal spinal reflexes and proper behavior in mice.

## INTRODUCTION

Smooth and flawless overground movement in mammals requires a timely and carefully orchestrated assembly and refinement of spinal sensory-motor circuits. Sensory neurons, spinal interneurons, and motor neurons are key players that form intricate neuronal circuits during embryonic development ^1^. These circuits are fine-tuned during early postnatal development, at a time where the animal is moving independently ^2–4^. Early formation of neuronal circuits is driven by specific molecular mechanisms ^5–7^. In sensory-motor circuits embedded within the spinal cord, information from the periphery, originating from proprioceptive and tactile sensory neurons, is essential in the generation of smooth movement and key to counteract perturbations during locomotion ^8–11^. Proprioceptors are a select set of sensory neurons that convey information about muscle stretch and tension, two metrics that enable the CNS to estimate limb kinetics and position ^12, 13^. Aside from supraspinal targets, proprioceptive neurons also feed into spinal reflex circuits through contacts with several classes of spinal interneurons ^14–17^ and direct monosysnaptic contacts onto motor neurons ^18, 19^.

In mature neuronal circuits, each motor neuron receives proprioceptive inputs that originate from the same (homonymous) muscle innervated by the motor neuron, as well as from proprioceptive fibers arising from synergistic muscles ^19, 20^. While mature motor neurons do not receive proprioceptive inputs from antagonistic muscles, during early development this is not the case. It has long been debated whether supernumerary sensory synapses are generated during spinal cord development, leading to formation of inappropriate contacts and irregular neuronal circuits ^16, 21–24^. Although it is currently unclear how and why these inappropriate synapses are formed, they are required to be eliminated for the emergence of mature sensory-motor circuits and proper motor function ^23, 24^. The molecular mechanisms responsible for the refinement of sensory-motor circuits and synaptic elimination remain poorly understood.

A subset of transcription factors (Hoxd9-11) has previously been implicated in irregular formation of select sensory-motor circuits, while other circuits are not affected ^25^. In addition, C1q, the initiating protein of the classical complement cascade, has been reported to tag synapses in several neuronal circuits, including visual system and cortical circuits ^26–28^. Furthermore, we have previously reported that C1q, is implicated in synaptic dysfunction and subsequent elimination during normal development and in the neurodegenerative disease spinal muscular atrophy ^29^.

An opposing scenario responsible for synaptic protection has implicated CD47, an integrin-associated protein, which has been described as a “don’t eat me” signal in the visual system ^30^. Furthermore, under pathological conditions, tumor cells overexpress CD47 to avoid engulfment by macrophages ^31^. In contrast, studies in cortical and hippocampal regions have reported that elimination of SIRPα, the CD47 receptor, contributes to synapse elimination at late but not early stages of development ^32^. Thus, the function of CD47 in synapse elimination is currently unclear.

To elucidate, the molecular mechanisms of synaptic elimination and their potential role in refinement of sensory-motor circuits during early development, we investigated whether activation of the classical complement cascade is causally involved. Specifically, we investigated the involvement of classical complement proteins, C1q and C3, in the spinal sensory-motor circuits during postnatal mouse development. We also studied whether CD47 is acting as a protecting agent of synapses. To address these questions, we used the well-established proprioceptive-motor neuron circuit responsible for the stretch reflex in the hindlimb of neonatal mice. Using virally-mediated neuronal map strategies, physiology, behavioral assays and mouse genetics to eliminate ubiquitously C1q, C3 and CD47, either alone or in combination, we investigated systematically the cascade of events involved in the formation and function of sensory-motor spinal circuits.

In both C3 and C1q knockout mice, we observed a robust increase of proprioceptive synapses within the motor neuron pools during early postnatal ages. Remarkably, we also observed a similar result in CD47 knockout mice, in contrast to its neuroprotective function in the brain. We determined that this aberrant increase (∼30%) of proprioceptive synapses in mutant mice, resulted in inappropriate synaptic contacts between flexor proprioceptors and antagonistic (extensor) motor neurons, and vice versa. These inappropriate synapses were functional and responsible for altered functional characteristics in motor neurons, leading to an abnormal behavioral phenotype. Taken together, our results identify two distinct molecular mechanisms - C3 through classical complement activation and CD47-SIRPα interactions - that mediate engulfment and removal of inappropriate synapses by microglia within spinal sensory-motor circuits.

## RESULTS

### Genetic deletion of C3 or CD47 results in supernumerary proprioceptive synapses

To investigate whether the classical complement pathway is involved in the elimination of supernumerary synapses in the developing spinal cord, we examined the role of C3 protein, a key regulator protein in the activation of the classical complement pathway ^33, 34^. At the same time, we investigated the potential protective role of CD47, since it has been reported to play a protective - “don’t eat me” - role for synapses in the retinogeniculate system ^30^.

To address the involvement of C3 and CD47 in synaptic regulation in the developing spinal cord, we assessed proprioceptive synapses (marked by VGluT1 and apposed on motor neurons) in the 4^th^ or 5^th^ lumbar (L4 or L5) spinal segments during early postnatal development in wild type (WT), C3 knock out (C3^-/-^), CD47 knock out (CD47^-/-^) and C3+CD47 double knock out (C3^-/-^CD47^-/-^) mice. The somatic and dendritic (0-50 μm distance from the soma) coverage of motor neurons (ChAT+ located in the ventral horn) by proprioceptive synapses was analyzed by immunohistochemistry and quantified using confocal microscopy at the postnatal ages, P1, P5 and P10 (Fig. 1A; Extended Data Fig. 1A-D). C3^-/-^CD47^-/-^ mice were viable only for a few days after birth (died ∼ P2/3). We found that the proprioceptive synaptic coverage increased with age in WT mice (blue bars in Fig. 1B,C). At P1, there was no significant difference between WT, C3^-/-^ and CD47^-/-^ mice. In contrast, C3^-/-^CD47^-/-^ mice revealed a significant higher incidence of synapses on both soma (Fig. 1B) and dendrites (Fig. 1C) of motor neurons compared to WT mice. Importantly, we observed a significantly higher incidence of proprioceptive synapses in C3^-/-^ mice at P5 and P10 compared with their age-matched counterparts (Fig. 1B,C). Remarkably, CD47^-/-^ mice also revealed a higher incidence of proprioceptive synapses, similar to that observed in C3^-/-^ mice. To investigate whether combined knock out of C3 and CD47 would result into added increase of synapses, we compared single and double heterozygous mice with their WT counterparts at P10, since double homozygous mice do not survive past P2/3. We found that C3^+/-^ (Extended Data Fig.1A,B) and CD47^+/-^ (Extended Data Fig.1A,C) mice possess significantly higher number of synapses in motor neuron somata at P10. Notably, combined heterozygous mice (C3^+/-^CD47^+/-^) revealed a significantly higher incidence of synapses compared to individual heterozygous mice (Extended Data Fig.1D). Together, these results demonstrate that supernumerary synapses form in the spinal cord during early development, and suggest that both classical complement C3-dependent and independent mechanisms are responsible for their elimination. Furthermore, it indicates that CD47 plays a surprisingly different and opposing role in the spinal cord compared to that in the brain and participate in synaptic removal.

**Figure 1.**
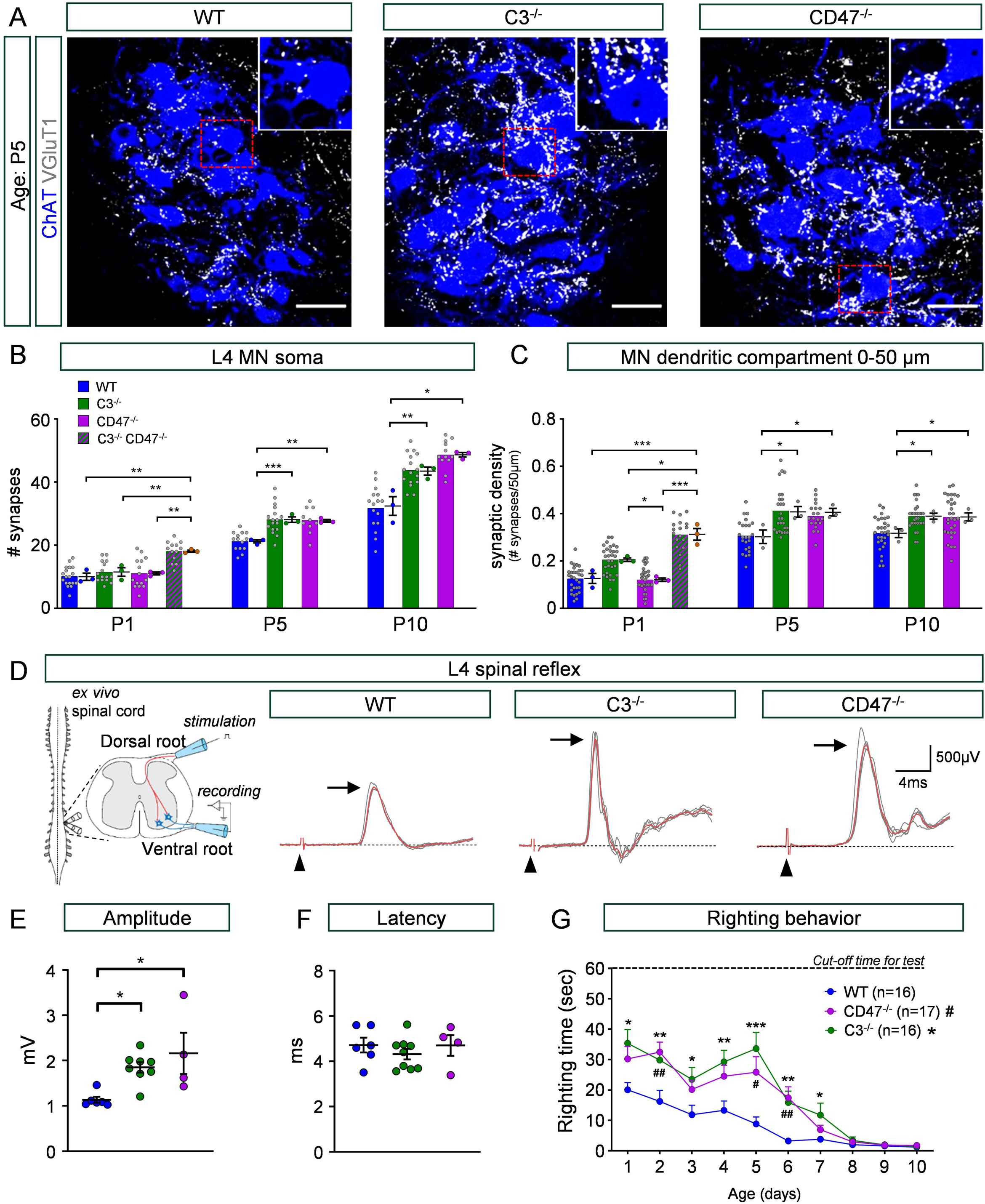
Genetic deletion of C3 or CD47 results in a higher incidence of proprioceptive (VGlut1+) synapses on spinal motor neurons. **A)** Confocal images of L4 motor neurons (ChAT, blue) and VGluT1 synapses (VGluT1, white) in WT, C3^-/-^ and CD47^-/-^ mice at P5. Insets show higher magnification images of a single motor neuron. Scale bar: 50 μm. **B)** Number of VGluT1 synapses per motor neuron soma at P1, P5 and P10 (N=3 mice/group; colored data points). Number of somata: [WT: P1, n=18; P5, n=14; P10, n=16] [C3^-/-^: P1, n=18; P5, n=17; P10 n=16] [CD47^-/-^: P1, n=15; P5, n=10; P10, n=12]. **C)** Synaptic density of proximal dendrites (0-50μm from the soma, N=3 mice/group). Number of dendrites: [WT: P1, n=30; P5, n=24; P10, n=30] [C3^-/-^: P1, n=30; P5, n=30; P10, n=30] [CD47^-/-^: P1, n=30; P5, n=20; P10, n=30]. Significance: *p < 0.05, **p < 0.001, ***p < 0.001; one-way ANOVA multiple comparisons with Bonferroni’s test. Statistical comparison was performed for average values in mice (colored data points). **D)** Simplified schematic of the *ex vivo* spinal cord preparation to assess the dorsal-ventral reflex at P5. Representative recordings of L4 ventral root responses following L4 dorsal root stimulation in WT, C3^-/-^ and CD47^-/-^ mice. Red trace is the average of the first five trials which are shown in grey. Arrowheads indicate stimulus artifacts. Black arrows indicate peak amplitude in the averaged response. **E)** Amplitude of spinal reflexes at P5. [WT: N=6 mice; C3^-/-^: N=8; CD47^-/-^: N=4]. Significance: *p < 0.05, one-way ANOVA multiple comparisons with Bonferroni’s test. **F)** Latency of responses for the same group as in (E). **G)** Righting times across the first ten postnatal days for WT, C3^-/-^ and CD47^-/-^ mice. Significance: (*) (#) p < 0.05, (**) (##) p < 0.01, (***) (###) p < 0.001 (compared to WT: CD47 significance #; C3 significance *), One-way ANOVA multiple comparisons with Bonferroni’s test.

We next sought to investigate whether the supernumerary proprioceptive synapses are functional in both C3^-/-^ and CD47^-/-^ mice. To address this, we performed physiological experiments in which proprioceptive synapses were tested using the *ex vivo* spinal cord. We quantified the amplitude of the 4^th^ lumbar (L4) dorsal-to-ventral root spinal reflex using the *ex vivo* spinal cord preparation (Fig. 1D) as we previously described ^29, 35^. We found that the L4 reflex amplitude was significantly increased in C3^-/-^ and CD47^-/-^ mice compared to their WT counterparts at P5 (Fig. 1D,E). The latency of the monosynaptic responses however, did not show any significant difference among the three experimental groups (Fig. 1F). These results demonstrate that the supernumerary synapses are functional in both C3^-/-^ and CD47^-/-^ mice. To test whether excessive proprioceptive synapses are susceptible to synaptic depression, we challenged these synapses at two different frequencies (0.1 and 10Hz). We found no differences in synaptic depression between the three groups, both at P5 and P10 (Extended Data Fig. 1E,F). This result indicates that supernumerary synapses do not differ in the neurotransmission reliability compared to WT counterparts.

To investigate the impact of excessive number of synapses on mouse behavior, we quantified the righting time in mice and found significantly longer latencies in both C3^-/-^ and CD47^-/-^ mice compared to WT controls in the first postnatal week (Fig. 1G; Video 1). To this end, heterozygous mice for C3 and CD47 also exhibited longer latencies to right (Extended Data Fig. 1I, Video 2), especially during the first five days after birth, where proprioception plays a major role in this behavior ^36^. In addition, heterozygous mice did not exhibit any significant body weight gain (Extended Data Fig. 1J), but comparison in homozygous mice for C3 or CD47 deletion exhibited some differences in body weight gain (Extended Data Fig. 1H). Taken together, these results indicate that both C3 and CD47 play a major role in the elimination of supernumerary synapses which in turn have significant effects on the mouse behavior.

### Supernumerary synapses are largely inappropriate and are eliminated via C3 or CD47 activity

Deletion of C3 or CD47, resulting in a higher incidence of proprioceptive synapses, raised the possibility that these synapses are inappropriate. Inappropriate synapses are considered those between sensory fibers originating from a flexor muscle and contacting an antagonistic motor neuron innervating an extensor muscle, and vice versa. If this hypothesis is correct, inappropriate synapses are destined to be eliminated during early development as part of a refinement process and the emergence of mature sensory-motor circuits. To distinguish between these possibilities, we examined whether proprioceptive fibers that innervate the dorsi-flexor muscle Tibialis Anterior (TA), make inappropriate contacts onto motor neurons that innervate the biomechanically antagonistic muscle gastrocnemius (Gs; plantar extensor muscle) and vice versa. To determine this, we utilized a virally-mediated mapping strategy combined with genetically modified mice, similar as described previously ^37^. Specifically, we used a PV:FlpO allele (targeting proprioceptors) ^38–41^, and the Cre and FlpO-dependent TdTomato reporter mice (Ai65:tdT) to label select proprioceptive fibers and their synapses following muscle injections of a virally Cre-expressed driver. To this end, PV::FlpO were crossed with the Ai65 Cre/FlpO dependent TdTomato reporter and the resulting animals (Pv::FlpO^+/+^; Ai65^+/+^) were crossed with C3^-/-^, C1q^-/-^, and C47^-/-^ mouse lines to obtain: *Pv::FlpO^+/+^/Ai65^+/+^*/*C3^-/-^*; *Pv::FlpO^+/+^*/*Ai65^+/+^/C1q^-/-^*; *Pv::FlpO^+/+^/Ai65^+/+^/CD47^-/-^* strains respectively.

To activate expression of the fluorescent tdTomato reporter in Gs proprioceptors, we injected a canine adeno virus 2 expressing Cre (abbreviated: CAV2^CRE^) ^42^ into the Gs muscle at P0. In addition, we simultaneously injected an rAAV6/ds-CBH-GFP (abbreviated: AAV6-GFP) virus in the TA muscle to drive expression of GFP and label the soma and dendrites of TA motor neurons (Fig. 2A,B). To validate this approach, we also injected both CAV2^CRE^ and AAV6-GFP into TA, to verify homonymous synapses between TdTomato, VGluT1+ proprioceptive fibers and synapses, and GFP TA motor neuron (ChAT) soma and dendrites in neonatal mice (Extended Data Fig. 2A).

**Figure 2.**
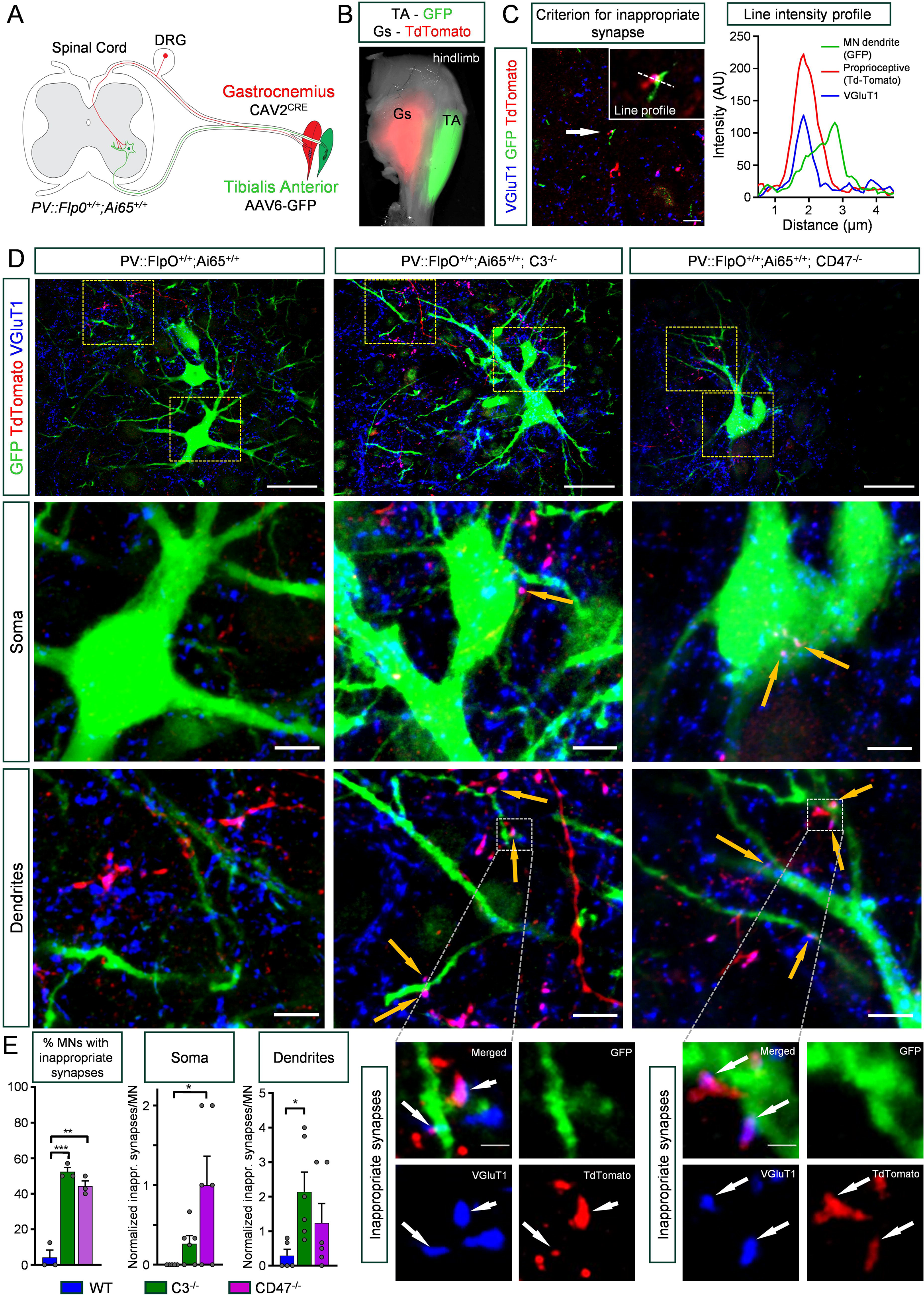
Inappropriate synapses on motor neurons formed by proprioceptive fibers originating in antagonistic muscles in C3^-/-^ and CD47^-/-^ mice. **A)** Schematic revealing the strategy for labelling proprioceptive fibers originating in gastrocnemius (Gs) muscle and labelling of motor neurons innervating the tibialis anterior (TA) muscle. TA motor neurons were labeled by injection of rAAV6/ds-CBH-GFP in TA muscle, and Gs proprioceptors were labeled by injection of CAV2^CRE^ virus in the Gs muscle in PV::FlpO;Ai65 mice. Injections were performed at P0/1 and tissue was harvested at P6/7. **B)** Superimposed fluorescent and transmitted-light photograph of the hindlimb injected at P5, showing Gs muscle in red (TdTomato) and TA muscle in green (GFP). **C)** Confocal image of an inappropriate synapse. Inset shows the line profile across the synapse (white arrow in low mag image) which was used as our criterion for synaptic contact. Graph shows the signal intensity for each of the three immunoreactive signals. Gs proprioceptive fibers labeled with DsRed (red), TA MN dendrites (green) and VGlut1 synapses (blue). **D)** Confocal images of L4 TA motor neurons from PV::FlpO;Ai65 (left), PV::FlpO;Ai65;C3^-/-^ (middle), and PV::FlpO;Ai65;CD47^-/-^ (right) mice. Z-stack distance: 3.5 μm. Top row, scale bar: 100 μm. Magnified image from yellow dotted box in the soma (center row) and dendrite (bottom row). Yellow arrows point to inappropriate synapses. Scale bar: 20 μm. Higher magnification insets from white dotted boxes showing inappropriate synapses on dendrites (bottom row) in PV::FlpO;Ai65;C3^-/-^ and PV::FlpO;Ai65;CD47^-/-^ mice only. White arrows highlight inappropriate synapses. Scale bar: 2 μm. **E)** Percentage of motor neurons with inappropriate synapses (both soma and dendrites). Left graph, WT: N=3; C3^-/-^: N=3; CD47^-/-^: N=3 mice (age: P5-P7). Middle and Right graphs: normalized number of inappropriate somatic (center) and dendritic (right) synapses per motor neuron (WT: N=5; C3^-/-^: N=6; CD47^-/-^: N=6 sections from 3 mice per group). Significance: *p < 0.05, **p < 0.001, ***p < 0.001; one-way ANOVA multiple comparisons with Bonferroni’s test.

Next, we investigated putative inappropriate synapses between Gs-originated proprioceptors and TA motor neurons at P6/7. Our criterion of an inappropriate synapse required colocalization of TdTomato in a Gs-proprioceptor fiber with VGluT1 immunoreactivity, which in turn, was apposed to a soma or dendrite of a TA motor neuron (labelled with GFP). This criterion was met through line intensity profiling (Fig. 2C) from single optical plane confocal images (see Methods). For quantification purposes, only VGluT1 (proprioceptive synaptic marker) and TdTomato (proprioceptive fiber/synapse originating from an identified muscle) signals partially overlapping with the GFP (motor neuron) signal were considered as inappropriate synapses (Fig. 2C). If the colocalized TdTomato and VGluT1 signals were not apposed to GFP (Extended Data Fig. 2B), these synapses were excluded from analysis. Under such stringent criteria, we found nearly complete absence of inappropriate synapses on either soma or proximal dendrites of TA motor neurons in WT mice (Fig. 2D,E). In striking contrast, ∼55% of TA motor neurons in C3^-/-^ and ∼45% of TA motor neurons in CD47^-/-^ mice exhibited a significantly higher number of inappropriate synapses (Fig. 2D,E). Additionally, we observed that the location of inappropriate synapses was different in each of the two mutant mouse lines. CD47^-/-^ mice revealed a significant increase in inappropriate synapses on the soma of TA motor neurons, while C3^-/-^ animals exhibited more inappropriate synapses on proximal dendrites (Fig. 2D,E). We also investigated whether there is a preferential direction of inappropriate proprioceptive synapses originating from dorsi-flexor muscles (i.e. TA muscle) to plantar extensor (i.e. Gastrocnemius) motor neurons. To address this possibility, we reversed the injections of the two viruses injected (CAV2^CRE^ in TA, and AAV6-GFP in Gastrocnemius). Our analysis revealed that inappropriate synapses were also formed between TA proprioceptors to Gs motor neurons in C3^-/-^ mice to a similar extent (Extended Data Fig. 2C,D). To provide further support that classical complement cascade is involved in synaptic refinement, we examined whether spinal motor neurons receive inappropriate synapses in C1q knockout mice. C1q is the initiating protein in the classical cascade pathway. As expected, C1q knock out mice, revealed ∼50% of motor neurons receiving inappropriate synapses (Extended Data Fig. 2E,F).

Lastly, we investigated whether the inappropriate synapses in C3^-/-^ and CD47^-/-^ mice might be the result of scrambled motor neuron position within the ventral horn, since previous reports demonstrated that when motor neuron position is altered, the proper organization of sensory-motor circuits is lost ^37, 43^. To address this possibility, TA and Gs motor neurons were selectively labelled with two different fluorescent tracers injected from their corresponding muscle (CTb488 in TA and CTb555 in Gs) at P0. At P5, spinal cord sections containing L4/L5 lumbar segments were scanned, mapped with Neurolucida (see Methods) and density maps were constructed indicating the dorso-ventral and medio-lateral extent of both TA and Gs motor neuron pools in WT, C3^-/-^ and CD47^-/-^ mice (Extended Data Fig. 2G). Using the central canal as a reference point, the contour of the transverse area in which TA and Gs motor neurons were positioned within the ventral horn according to the coordinates was identified, mapped and statistically compared. We found no significant differences in the density map of the location for either TA motor neurons (Extended Data Fig. 2H) or Gs motor neurons (Extended Data Fig. 2I; Video 3) in all groups.

Taken together, these results demonstrate that excessive synapses between proprioceptors and antagonistic motor neurons are formed during early development, resulting in miswired sensory-motor circuits and this miswiring is not attributed to the location of motor neurons.

### Inappropriate sensory synapses are functional, resulting in improper muscle contraction

We next investigated whether the inappropriate synapses are functional by performing H-reflex experiments using the *ex vivo* spinal cord-hindlimb preparation ^15, 44^ at P5 (Fig. 3A). The H-reflex (or Hoffman reflex) is the reaction in muscles following electrical stimulation of proprioceptive sensory fibers within their innervating nerves through the spinal cord (Fig. 3A) ^45, 46^. The M-response is the muscle response following direct stimulation of motor neurons axons within the stimulating nerve. In WT mice, stimulation of the common peroneal nerve resulted in a robust M-response and an H-response (or H-reflex) in the TA muscle (Fig. 3B). In addition, simultaneous recordings from the antagonistic muscle Gs did not result in any responses, as expected in healthy (WT) control mice (Fig. 3B). We confirmed that the neurotransmitter responsible for the production of the H-reflex was glutamate, since exposure to the NMDA receptor blocker D-AP5 (50μM) and the AMPA/kainate receptor blocker NBQX (20μM), abolished the response leaving relatively unaffected the M-response (Fig. 3B, right traces). In C3^-/-^ and CD47^-/-^ mutant mice however, a response with a latency similar (or slightly delayed) to the H-reflex was detected in the antagonistic Gs muscle, following CP nerve stimulation (Fig. 3C; and insets). The amplitude of the H-reflex induced in the homonymous muscle (CP nerve stimulation to TA muscle) did not reveal any significant differences between the three experimental groups (Fig. 3D in CP→TA). In contrast, we observed an inappropriate response in the antagonistic Gs muscle, with similar latency to that of the H-reflex observed in the homonymous TA muscle (Fig. 3C, D in CP→Gs, and E), only in C3^-/-^ and CD47^-/-^ mice. This result suggests that inappropriate synapses derived from TA muscle afferents, are sufficient to activate Gs motor neurons, resulting in improper Gs muscle contraction (Fig. 3C, blue arrows). Similar results were obtained when we tested stimulation of tibial nerve and recorded in the TA muscle (Fig. 3F,G).

**Figure 3.**
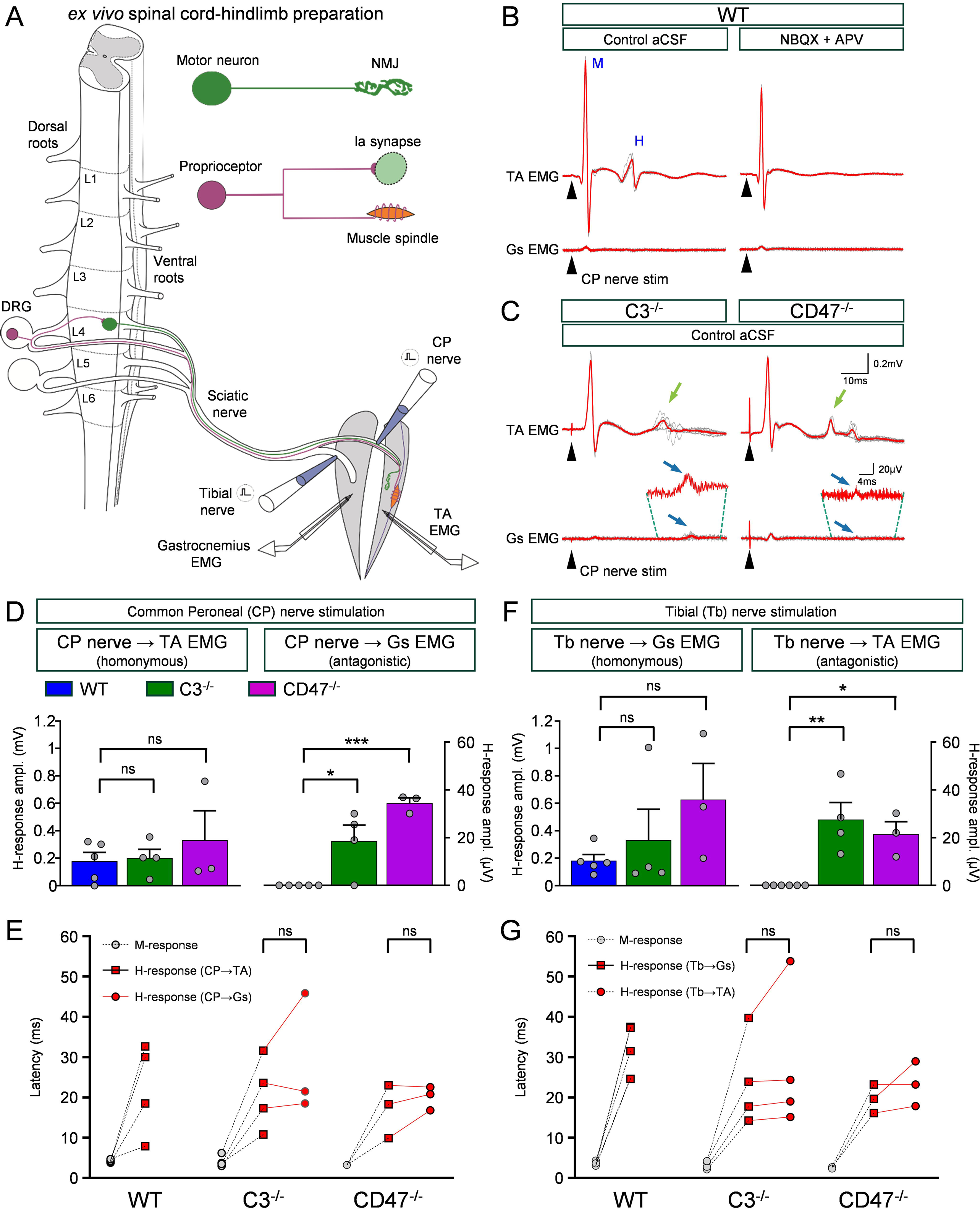
Inappropriate proprioceptive sensory synapses on motor neurons induce responses in antagonistic muscles. **A)** Schematic of the *ex vivo* spinal cord-hindlimb preparation. Suction electrodes were placed on the common peroneal (CP) and tibial (Tb) nerves to perform “*en passant*” stimulation. Bipolar concentric needle electrodes recorded EMG activity from the TA and Gs muscles. A motor neuron and a neuromuscular junction (NMJ) are shown in green. A proprioceptor with its muscle spindle and the Ia central synapse on motor neurons are shown in purple. **B)** Simultaneous EMG recordings from TA and Gs muscles following CP nerve stimulation, in a WT spinal cord under control aCSF solution, and after exposure to NBQX (20 μM) and D-AP5 (50 μM). Red traces show the average of five trials (grey). Black arrowhead denotes stimulus artifact. **C)** Traces from C3^-/-^ (left) and CD47^-/-^ (right) mice in control aCSF. Green arrows point to homonymous muscle H-reflex, while blue arrows point to inappropriate H-responses from the antagonistic Gs muscle following CP nerve stimulation. **D)** Amplitude of H-reflex induced by CP nerve stimulation in the homonymous TA muscle (left; CP nerve → TA EMG), and the H-response from the antagonistic Gs muscle (right; CP nerve → Gs EMG). **E)** Latency measurements of M-responses evoked by CP nerve stimulation in the TA muscle (grey circles), as well as H-reflex in TA muscle following CP nerve stimulation (red squares), and antagonistic H-responses in the Gs muscle following CP nerve stimulation (red circles). **F)** Amplitude of H-reflex induced by Tb nerve stimulation in the homonymous Gs muscle (left; Tb nerve → Gs EMG), and the H-response from the antagonistic TA muscle (right; Tb nerve → TA EMG). Each data point corresponds to a single animal. [WT: N=5; C3^-/-^: N=4; CD47^-/-^: N=3 mice]. Significance: *p < 0.05, **p < 0.01, ***p < 0.001; One-way ANOVA multiple comparisons with Bonferroni’s test. ns: no significance. **G)** Latency measurements of M-responses evoked by Tb nerve stimulation in the Gs muscle (grey circles), as well as H-reflex in Gs muscle following Tb nerve stimulation (red squares), and antagonistic H-responses in the TA muscle following Tb nerve stimulation (red circles). One-way ANOVA multiple comparisons with Bonferroni’s test; ns: no significance.

We next investigated the hypothesis that if the classical complement cascade is involved in refinement of inappropriate synapses, C1q^-/-^ mice are expected to exhibit similar inappropriate responses to C3^-/-^ mice. To this end, our experiments using C1q^-/-^ mice confirmed resulted in inappropriate H-reflex responses in the Gs muscle following stimulation of the antagonistic CP nerve (Extended Data Fig. 3A-C) validating our hypothesis. These results demonstrate that both C3 and CD47 proteins are involved in the pruning of supernumerary synapses, formed during embryonic and early postnatal development.

We also wanted to exclude the possibility that the inappropriate responses in the antagonistic muscles were F-waves ^47^. However, repeated stimulation of the nerve at low frequencies (∼0.1Hz) revealed that the responses were time locked (F-waves are not time locked) and appeared earlier than the typical F-wave responses which were variable in amplitude and latency (Extended Data Fig. 3D). The latency of the inappropriately-evoked EMG response in the Gs muscle after CP nerve stimulation varied considerably (range: -1ms to +15ms; Fig.3 E,G) raising the possibility that inappropriate proprioceptive synapses might not be synapsing directly on motor neurons. To resolve this, we performed whole cell patch clamp recordings using the *ex vivo* spinal cord-hindlimb preparation to record intracellular responses from TA motor neurons following selective peripheral nerve stimulation (Fig. 4A). The ventral roots L4 and L5 were cut to avoid contamination of synaptic currents by the antidromic action potential induced by the homonymous (CP) nerve stimulation. Both L4 and L5 ventral roots were placed in suction electrodes, so the patched neuron could be identified as a motor neuron by the presence of an antidromic action potential (Extended Data Fig. 4 A,B). TA motor neurons were visually targeted for whole cell intracellular recordings by the presence of CTb-488, which was previously injected into the TA muscle at P0 (Extended Data Fig. 4E). Excitatory postsynaptic currents (EPSCs) induced by either CP nerve (homonymous) or tibial nerve (antagonistic) stimulation were compared in WT, C3^-/-^ and CD47^-/-^ mice. To assess if the responses were monosynaptic in nature, we analyzed the latency of the EPSCs at different frequency of stimulation (0.1, 0.2 and 1Hz) and found that the coefficient of variation (jitter test) was not significantly different (Extended Data Fig. 5A-C), demonstrating monosynapticity. Our criterion for the duration of monosynaptic transmission in our neonatal *ex vivo* spinal cord was 3.5ms, as we previously reported ^20^.

**Figure 4.**
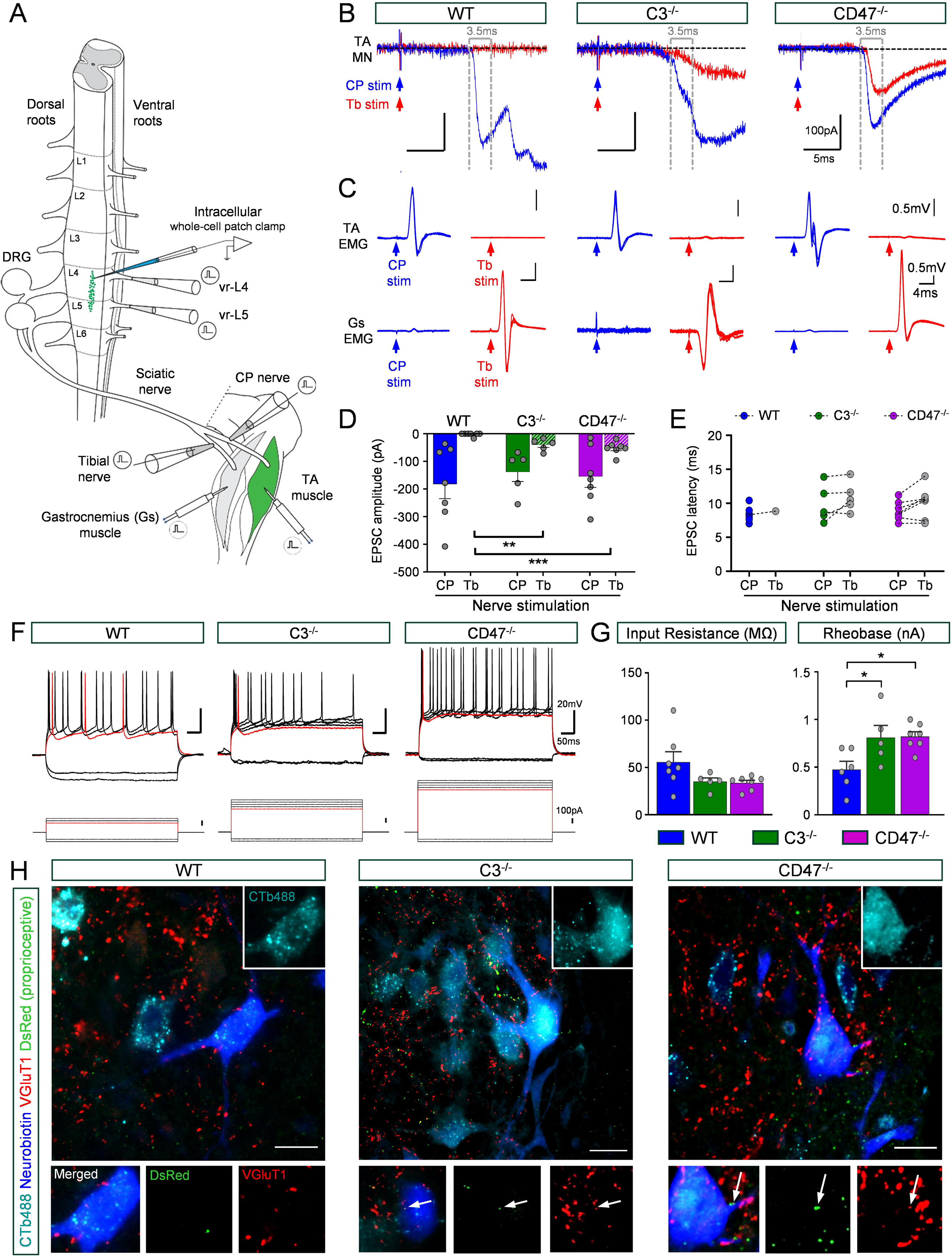
Inappropriate synapses evoke monosynaptic EPSCs. **A)** Schematic of the *ex vivo* spinal cord-hindlimb preparation utilized for whole cell patch-clamp from TA motor neurons. Bipolar concentric needle electrodes were placed in the TA and Gs muscles to record EMG. Suction electrodes were placed in the CP and Tibial nerves for stimulation “*en passant*”. The L4 and L5 ventral roots were cut and placed in suction electrodes. Intracellular electrode contained Neurobiotin. At birth, the TA muscle was injected with CTb-488, while the Gs muscle was injected with CAV2^CRE^ virus in PV::FlpO;Ai65-TdTomato mice. **B)** Whole-cell responses from TA motor neurons following stimulation of the homonymous CP nerve (blue) or the antagonistic Tb nerve (red) in WT, C3^-/-^ and CD47^-/-^ mice at P4-6. The EPSC response is the mean of five responses (evoked at 0.1Hz). Dotted lines depict the duration of 3.5ms, which was used as the criterion for monosynaptic duration. **C)** EMG responses corresponding to the experimental groups as shown in (B) above. The TA (top traces) and Gs EMG responses (bottom traces) were acquired concurrently following stimulation of either: the CP nerve (blue) or the Tb nerve (red). Superimposed traces shown are five responses evoked at 0.1 Hz. **D)** Amplitude of EPSC evoked in TA motor neurons following CP (plain bars) or Tb (hatched bars) nerve stimulation in WT (blue), C3^-/-^ (green) and CD47^-/-^ (purple) mice [WT: n=7 motor neurons; C3^-/-^: n=5; CD47^-/-^: n=7]. One motor neuron per mouse was recorded. **E)** Latency of EPSC for experimental groups shown in (D). Note that in WT mice, only one TA motor neuron exhibited a small EPSC following Tb nerve stimulation. **F)** Voltage responses in TA motor neurons following steps of current injection (in patch-clamp experiments) in WT, C3^-/-^ and CD47^-/-^ mice (as shown in B). **G)** Input resistance (left) and rheobase (right) of TA motor neurons in the three experimental groups. Significance: * p < 0.05, ** p < 0.01, *** p < 0.001; one-way ANOVA multiple comparisons with Bonferroni’s test. **H)** Confocal images of recorded, filled and subsequently visualized TA motor neurons from the three experimental groups: PV::FlpO;Ai65 (abbreviated: WT), PV::FlpO;Ai65;C3^-/-^ (abbreviated: C3^-/-^), and PV::FlpO;Ai65;CD47^-/-^ (abbreviated: CD47^-/-^) mice. Images correspond to TA recorded motor neurons for each mouse line from experiments shown in B, C and F. Quadruple immunostaining against VGluT1 (red), DsRed (green), Neurobiotin (blue), and injected CTb488 (magenta). Images are Z-stack projections of single optical planes totaling 3.5μm in the Z-axis. Inserts are single optical plane, corresponding to higher magnification of the recorded motor neuron soma (top; CTb); white arrow indicates inappropriate synapses on the soma of the TA motor neuron in C3^-/-^ and CD47^-/-^ mice only (bottom images). Scale bar 20 μm.

Stimulation of the homonymous nerve (CP) resulted in robust monosynaptic EPSCs in TA motor neurons in all experimental groups (blue traces in Fig. 4B) with a similar amplitude on average (Fig. 4D). In striking contrast, stimulation of the antagonistic nerve (tibial) resulted in monosynaptic EPSCs only in C3^-/-^ and CD47^-/-^ mice, but not in their WT counterparts (red traces in Fig. 4B). This indicates that responses evoked 3.5ms after the onset of the EPSC, are considered polysynaptic in nature. Importantly, the resultant EPSC latency revealed no significant differences for either CP or tibial nerve stimulation (Fig. 4E). Appropriate stimulation of each peripheral nerve was verified by EMG responses recorded in both TA and Gs muscles simultaneously (Fig. 4C). The Compound Muscle Action Potential (CMAP) latency was not significant different in the homonymous muscle (CP→TA or Tb→Gs) in all three groups (Extended Data Fig. 4C). This data demonstrate that inappropriate proprioceptive synapses result in monosynaptically-induced EPSCs in C3^-/-^ and CD47^-/-^ mice.

We next investigated the effects of inappropriate synapses on the active and passive membrane characteristics of TA motor neurons. We found that resting membrane potential in all TA motor neurons was not statistically different across the three groups (Extended Data Fig. 4B). The input resistance revealed a trend towards a reduction, while the rheobase current was significantly higher in both mutant mice compared to WT controls (Fig. 4F,G). The moderate increase in rheobase may underlie the reason for the trend in increase of the repetitive firing ability in C3^-/-^ and CD47^-/-^ motor neurons (Extended Data Fig. 4D).

To reveal the site of the inappropriate synapses on recorded TA motor neurons, we labelled motor neurons with Neurobiotin (added in the intracellular solution), and visually revealed *post hoc* (blue in Fig. 4H and Extended Data Fig. 4E). Proprioceptive fibers originating in Gs muscles were labelled using our strategy explained previously (Fig. 2A), in which CAV2^CRE^ was injected in Gs muscle. TA motor neurons were marked by CTb-488 (cyan in Fig. 4H and green in Extended Data Fig. 4E) injected in the TA muscle. In this assay, we found that inappropriate synapses were located on the soma and proximal dendrites of TA motor neurons, only in C3^-/-^ and CD47^-/-^ mice (white arrows in Fig. 4H) but importantly, not in WT mice. Taken together, these results demonstrate that inappropriate synapses on TA motor neurons are functional and alter their intrinsic properties, likely impacting muscle contraction.

### Expression of CD47 and its receptor SIRPα, in the spinal cord

We next sought to identify the source of CD47 in the spinal cord. We performed RNAscope experiments combined with immunohistochemistry assays in neonatal animals (P0-P10). We found that CD47 was expressed in motor neurons (identified by ChAT immunoreactivity) (Fig. 5A), in spinal interneurons in the intermediate grey matter (identified by NeuN antibodies in the ventral horn) (Fig. 5B) and in proprioceptive neurons (identified by parvalbumin immunoreactivity) (Fig. 5C). This is in partial agreement compared to a recent report in which CD47 was only expressed in postsynaptic, but not presynaptic, sites in the brain ^48^, unlike our study, where we observe CD47 both presynaptic (proprioceptor) and postsynaptic (motor neuron) sites. CD47 was not expressed in either microglia (identified by Iba1 immunoreactivity) or astrocytes (identified by GFAP immunoreactivity; Fig. 5A,B). Importantly, the CD47 RNAscope probe was validated in CD47^-/-^ spinal cord, in which no signal was detected (Fig. 5B; bottom row images). SIRPα - the CD47 receptor - was expressed exclusively in proprioceptors (Fig. 5D) and microglia (Fig. 5E,F), but not in motor neurons (Fig. 5E).

**Figure 5.**
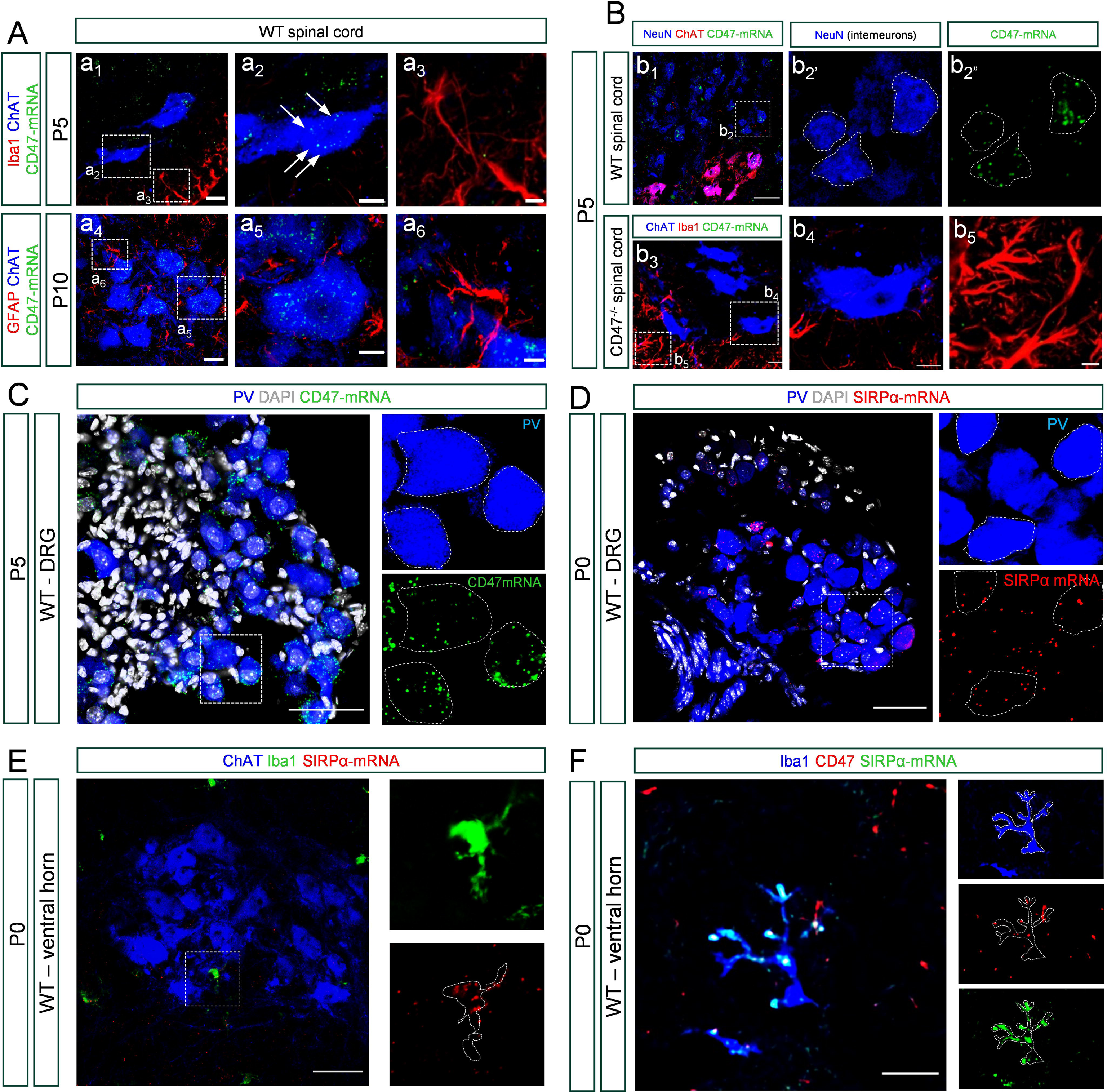
Neurons express CD47, while microglia express SRIPα. **A)** Z-stack confocal images (total distance in Z-axis: 3.5μm) from a P5 L4 spinal cord showing motor neurons (ChAT; blue), microglia (Iba1; red), and CD47-mRNA (RNAscope, in green) at P5 (top row). At P10 (bottom row), images show motor neurons (ChAT; blue), astrocytes (GFAP, red), and CD47-mRNA (green). Scale bar in a_1_ and a_4_: 10μm. Middle and right column correspond to higher magnification images marked by dotted boxes in a_1_ and a_4_ respectively. White arrows indicate the CD47-mRNA puncta. Scale bar in a_2_, a_3_, a_5_, a_6_: 5μm. **B)** Confocal images of CD47-mRNA puncta in interneurons, labelled by NeuN (blue), motor neurons (ChAT; red), and CD47-mRNA (green). Scale bar in b_1_: 50 μm. Bottom row of images show validation of CD47-mRNA specificity using an RNAscope assay in CD47^-/-^ mouse, with immunohistochemistry against ChAT (b_4_) and Iba1 (b_5_). Scale bar in b_3_, b_4_, b_5_: 10 μm. **C)** Confocal images of dorsal root ganglia (DRG) showing immunoreactivity against Parvalbumin (blue), DAPI signal (white) and RNAscope probe against CD47-mRNA (green). Scale bar 50 μm. **D)** Confocal image of Parvalbumin (blue), DAPI (white) and SIRPa-mRNA (red). Scale bar 50 μm. **E)** Confocal image showing motor neurons (ChAT, blue), microglia (Iba1, green) and SIRPα-mRNA (red). Scale bar 50 μm. Insets on the right, show higher magnification of Iba1 and SIRPα-mRNA in dotted box area. **F)** Single optical plane of confocal image showing immunoreactivity for microglia (Iba1, blue), and CD47 (red), and RNAscope signal against SIRPα-mRNA (green). Scale bar 5 μm.

### CD47 tags proprioceptive synapses on motor neurons

We next investigated the potential molecular mechanism involved in synaptic pruning and regulated by CD47 activity. CD47 has been described as the main molecule involved in synapse protection in the brain. However, in complete contrast, the results of our study demonstrate that CD47 is involved in synaptic elimination, an unanticipated function for CD47. To dwell deeper into this function of CD47, we investigated whether CD47 expression was involved in synaptic tagging and their engulfment by microglia cells in the developing spinal cord.

We performed immunohistochemistry experiments in which motor neurons (ChAT+) and their proprioceptive synapses (VGluT1+) were investigated for their potential association with CD47 at P1, P5 and P10, in WT, C3^-/-^ and CD47^-/-^ mice using a validated anti-CD47 antibody (Fig. 6 A). CD47 was considered tagging VGluT1+ synapses (apposed on either soma or dendrites of motor neurons) when there was an overlap of their corresponding immunoreactive optical signals (white arrows and graphs in Fig. 6 B,C). The number of CD47-tagged VGluT1+ synapses was significantly increased at P5 and P10 in C3^-/-^ mice, but not at P1 (Fig. 6 A,D,E) compared to WT mice. CD47 antibody specificity was validated in CD47^-/-^ mice (Fig. 6 A,D,E). CD47 increased synaptic tagging was also evident in C1q^-/-^ proprioceptive (VGluT1+) synapses on motor neurons, both on their soma and dendrites from P1 until P10 (Extended Data Fig. 6 A,B,C). To this end, genetic knock out of C1q ubiquitously resulted in a higher incidence of proprioceptive synapses on motor neurons from P5 onwards (Extended Data Fig. 6 D,E,F), in agreement with our previous report that the classical complement cascade is involved in synaptic elimination ^29^. Furthermore, the amplitude of the monosynaptic responses in C1q^-/-^ mice was also significantly higher compared to age-matched WT controls at P5 (Extended Data Fig. 6 G,H).

**Figure 6.**
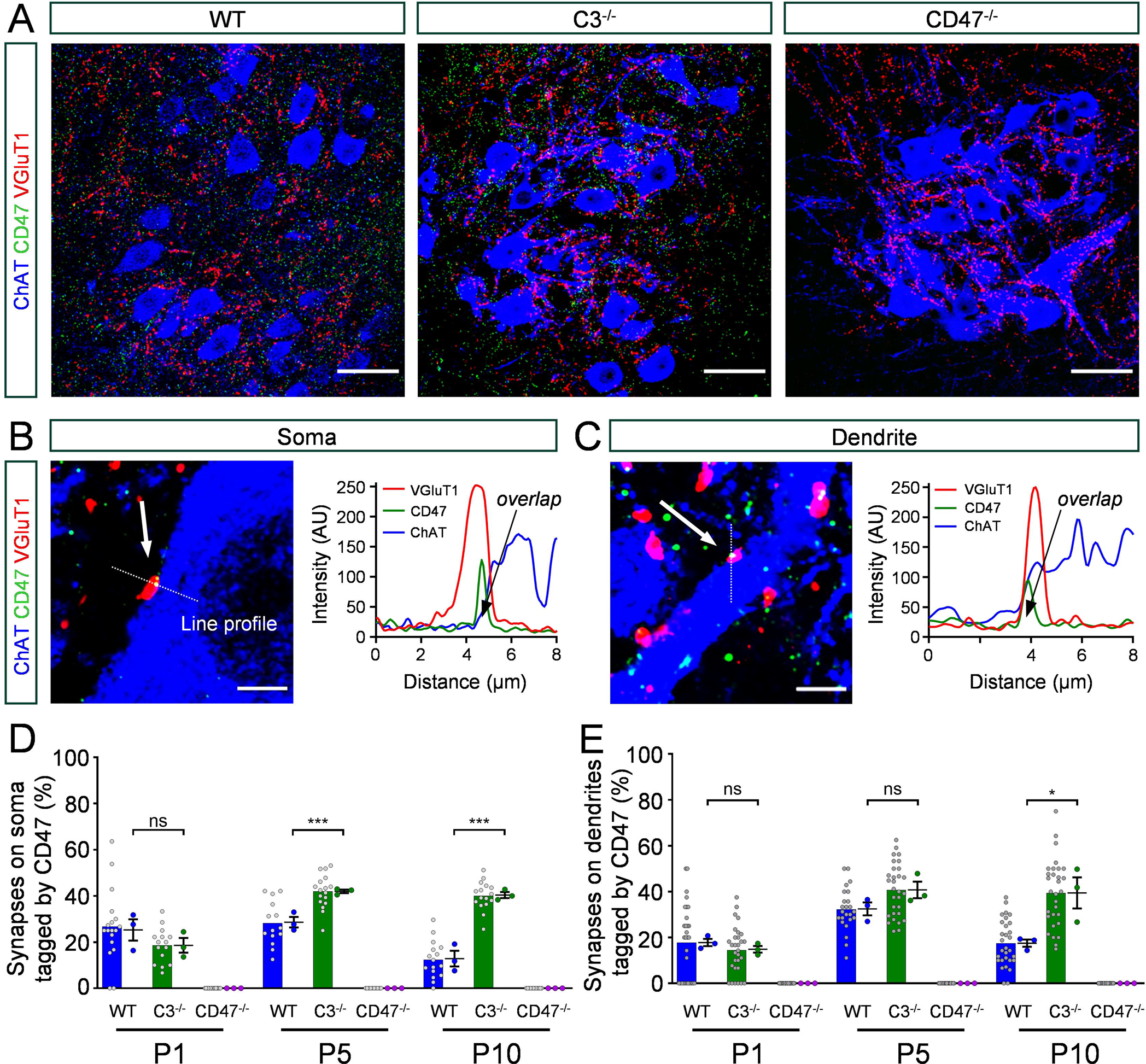
CD47 tags proprioceptive synapses in spinal sensory-motor circuits. **A)** Single plane confocal images of VGluT1 (red), CD47 (green) and ChAT (blue) immunoreactivity in WT, C3^-/-^ and CD47^-/-^ mice at P5. Scale bar: 50 μm. **B)** VGluT1+ synapse (red) apposed to a motor neuron soma (ChAT, blue) is tagged by CD47 (green). Graphs shows that the CD47 signal intensity overlaps with that of VGluT1 indicated by line profile (arrow, dotted line) for the three antibodies. Scale bar: 5 μm. **C)** Similar to (B) but for a synapse on a motor neuron dendrite. Scale bar: 5 μm. **D)** Percentage of somatic synapses tagged by CD47 at P1, P5 and P10. **E)** Percentage of dendritic synapses tagged by CD47. Each dot (in grey) corresponds to one motor neuron. Colored data points show average values from a single mouse (N=3 mice/group). # of motor neurons analyzed from N=3 mice [WT: P1, n=30; P5, n=24; P10, n=30] [C3^-/-^: P1, n=30; P5, n=30; P10, n=30] [CD47^-/-^: P1, n=30; P5, n=20; P10, n=30]. Statistical comparison was performed across average from mice. Significance: ***p < 0.001; one-way ANOVA multiple comparisons with Bonferroni’s test; ns: no significance.

Since CD47-tagging was higher in C3^-/-^ and C1q^-/-^ mice, we investigated whether CD47 might be upregulated in motor neurons. However, we observed no differences in CD47 mRNA puncta in TA motor neurons from C3^-/-^ mice (Extended Data Fig. 7 A,B) suggesting that CD47 tagging was unlikely to originate from motor neurons. In addition, there was no CD47 mRNA expression in either microglia or astrocytes in C3^-/-^ mice, similar to their WT counterparts (Extended Data Fig. 7 A,B). These results indicate that CD47 tagging of proprioceptive synapses is independent of the classical complement cascade. Furthermore, synaptic elimination would likely result as a co-operation between activation of the classical complement and CD47-mediated molecular mechanisms.

### VGluT1 synaptic remnants tagged with either CD47 only, or combined with C1q, are engulfed by microglia

Since we observed a higher CD47 tagging in either C3^-/-^ or C1q^-/-^ mice, we next investigated whether CD47 and classical complement are mutually exclusive for synaptic elimination. To test this, we performed immunohistochemistry for C1q and CD47 in the spinal cord of neonatal (P0-P5) mice. Remarkably, we found that both CD47 and C1q can tag the same VGluT1 synapse (yellow circles in Fig. 7 A,B). We also found synapses that were tagged either by CD47 or C1q alone (red circles for C1q and green circles for CD47 in Fig. 7 A,B). This synaptic tagging was increased as animals aged from P0 to P5 (Fig. 7 C).

**Figure 7.**
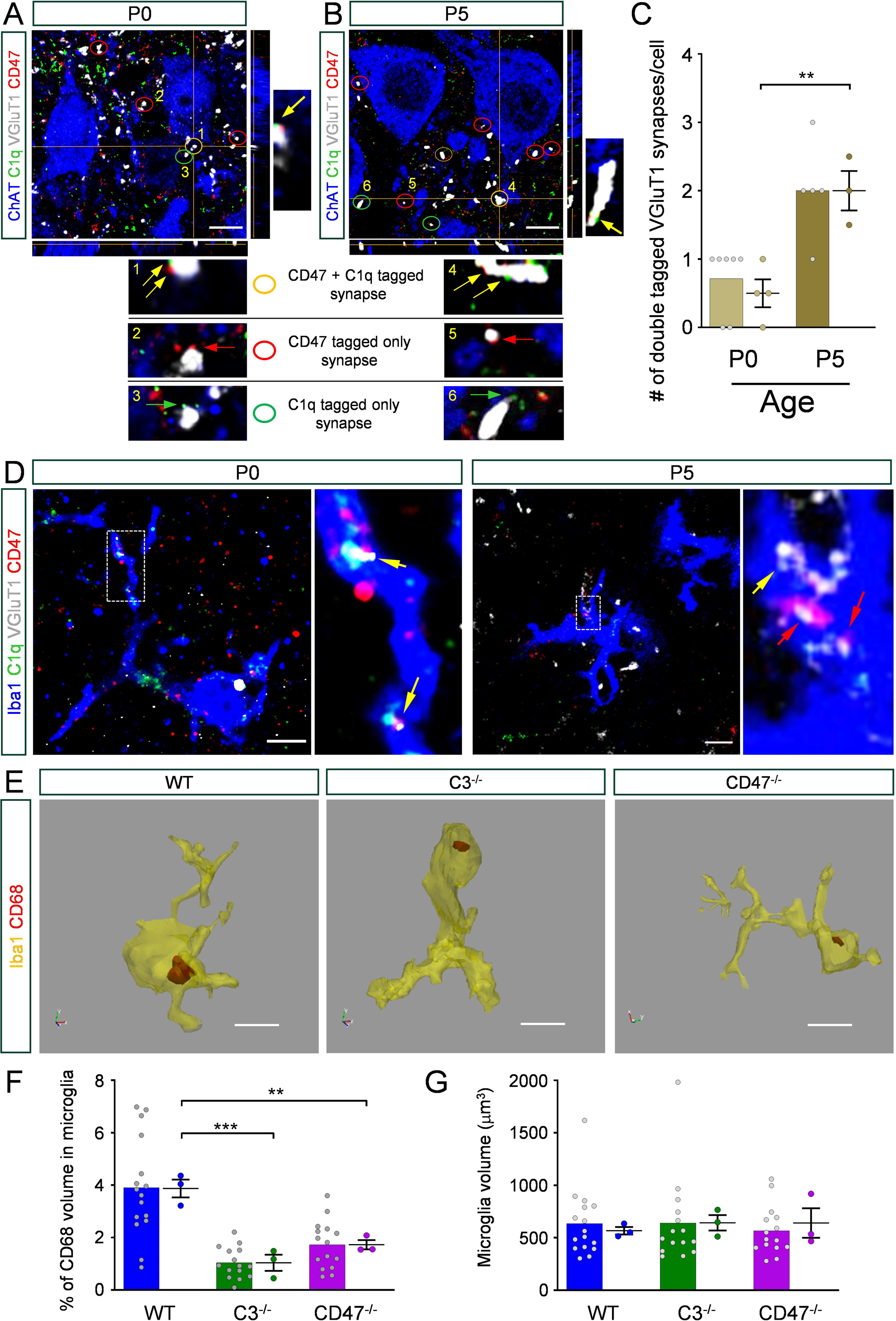
Microglia engulf synapses tagged by C1q, CD47 or combined. Orthogonal view of images immunoreactive against C1q (green), CD47 (red), VGlut1 (white) and ChAT (blue) at P0 **(A)** and P5 **(B)**. Circles indicate instances where a VGlut1 synapse is tagged by CD47 only (red circle), C1q only (green circle) or combined by both CD47 and C1q (yellow circle). Inserts at the bottom represent higher magnifications. Yellow arrows indicate double tagging of VGluT1 synapses by CD47 and C1q. Red arrows indicate tagging by CD47 only. Green arrows indicate tagging by C1q only. Scale bar in A and B: 10 μm. **C)** Quantification of VGluT1 synapses double-tagged at P0 (grey bar) and P5 (blue bar). Grey data points are number of synapses and colored data points are averages from single mice (N=4 at P0 and N=3 at P5). Significance: ** p < 0.01; Unpaired t-test. **D)** Single optical plane confocal images of immunoreactivity against Iba1 (blue), C1q (green), CD47 (red) and VGluT1 (white), at P0 and P5. Inserts, higher magnification of area indicated by dotted box. Yellow arrows indicate VGlut1 residues within microglia tagged by C1q and CD47. Red arrows indicate VGluT1 residues within microglia tagged by CD47 only. Scale bar: 5 μm. **E)** 3D reconstructions of microglia (Iba1; yellow) and CD68 (red), showing the presence of CD68 within microglia. **F)** Percentage of microglia containing CD68. Grey data points represent the number of microglia analyzed. Colored data points is the average from a single mouse (N=3 mice/group). **G)** The volume of microglia in WT, C3^-/-^ and CD47^-/-^ mice. Grey points are the number of microglia analyzed and colored data points are the average microglia volume per mouse (N=3 mice/group).

Since synapses tagged by C1q are susceptible to being engulfed by microglia ^28, 29^, we next investigated the possibility that synapses tagged by C1q and CD47 could also be eliminated in a microglia dependent manner. If CD47 played a protective role in synapse elimination, as reported in brain regions ^30^, by extension, it should prevent engulfment by microglia. In such case, it is expected that no CD47 should be found within microglia profiles. In contrast to this hypothesis, we found microglia containing VGluT1+ synaptic remnants tagged by both CD47 and C1q, at both P0 and P5 (yellow arrows in Fig. 7 D). Remarkably, we also identified VGluT1+ residues associated with only CD47 inside microglia, especially at P5 (red arrows in Fig. 7 D). These results suggest that microglia engulf synapses via CD47 tagging, in parallel to a mechanism requiring tagging by both CD47 and C1q. It is therefore likely that both of these mechanisms operate independent of each other.

Importantly, we identified interaction sites between SIRPα, CD47 and proprioceptive (PV+) synapses within the motor neuron pool in postnatal spinal cords (Extended Data Fig. 7 C). In further support, we observed SIRPα and CD47 in close association around the motor neuron (ChAT+) membrane (Extended Data Fig. 7 D,E), suggesting putative protein-protein associations with synapses impinging on motor neurons. This finding strongly suggests that a novel mechanism of synaptic engulfment by microglia is mediated by CD47 and its receptor SIRPα.

To provide further support of the involvement of microglia in synaptic elimination mediated by CD47, we investigated protein expression of CD68, a protein previously reported to correlate with engulfment activity ^34^. Using immunohistochemistry against Iba1 and CD68, confocal microscopy and Neurolocida, we measured the volume of CD68 within microglia in WT, C3^-/-^ and CD47^-/-^ mice (Fig 7 E). We found a significant reduction in the percentage of CD68 presence in microglia in C3^-/-^ and CD47^-/-^ mice compared to WT counterparts (Fig. 7 F), suggesting less synaptic engulfment in either C3 or CD47 knock out mice. In addition, there was no difference in the volume of microglia analyzed in any of the experimental groups (Fig. 7 G), indicating that activation of microglia in not altered across the three different groups.

Taken together, these results demonstrate that complement cascade is not the only source of synaptic elimination during normal development, but importantly, we implicate a previously unknown and unforeseen role for CD47, which instead of synapse protection, it contributes to synaptic pruning. Furthermore, the classical complement and CD47-mediated molecular mechanisms are not mutually exclusive and could operate in a co-operated manner. This liaison ensures refinement of sensory-motor circuits during early spinal cord development presented by a dual fail-safe mechanism.

## DISCUSSION

The understanding of the role of supernumerary synapses in spinal cord has been elusive, unlike that in the brain. Here we demonstrate that during early development, there is an excessive number of proprioceptive sensory synapses impinging on motor neurons. These extra synapses make inappropriate contacts with motor neurons, resulting in miswired immature sensory-motor circuits. The emergence of mature sensory-motor circuits is regulated by two molecularly distinct but yet, cooperating mechanisms. The first employs activation of the classical complement cascade, through C1q-C3 pathway, while the second utilizes CD47-SIRPα signaling. The involvement of CD47 in synaptic elimination within spinal cord is a totally unexpected function and in complete contradiction to the “don’t eat me” function in brain regions. Taken together, our study proposes that the natural course of elimination of inappropriately-generated synapses during early spinal cord development in mice, employs a dual fail-safe system to ensure the emergence of mature spinal reflexes and normal motor control occurs unimpeded.

### Immature spinal sensory-motor circuits are miswired during early development

We report that genetic removal of the classical complement proteins, C1q and C3, as well as the integrin associated protein CD47, result in ∼30% higher incidence of sensory synapses within the motor neuron pools in the developing lumbar spinal cord compared to normal mice. Our observations demonstrate that supernumerary synapses are established during embryonic development and subsequently pruned during postnatal development. The origin of these supernumerary synapses are proprioceptive. Despite previous reports of transient inappropriate sensory synapses ^20, 23, 29^, their precise origin and role are not understood. Here, we establish that proprioceptive sensory neurons make inappropriate synaptic contacts with motor neurons innervating a biomechanically antagonistic muscle. Supernumerary synapses are likely to be inappropriate synapses only, and not excessively-generated appropriate synapses, since the amplitude of H-reflex in the homonymous muscle between wild type, C3^-/-^ and CD47^-/-^ mice was similar. In addition, the origin of inappropriate synapses could be from several antagonistic muscles which are biomechanically similar in function (i.e. Tibialis Anterior and Extensor Digitorum Longus) ^20^.

It has been previously reported that miswired neuronal circuits may be the result of mislocated postsynaptic sites. In the case of a spinal sensory-motor circuit, the *FoxP1* gene can control the spatial and settling position of motor neurons, leading non-specific sensory-motor circuits ^43^. However, our analysis in the tibialis anterior and gastrocnemius motor neuron pools did not reveal any significant differences in the final settling position of their motor neuron somata when C3 or CD47 were genetically eliminated ubiquitously in mice. Importantly, we show that the inappropriate synapses are functional and exert an influence on the intrinsic firing properties of motor neurons they impinge upon. This miswired and immature circuit results in improper muscle contraction, and negatively impacts initial development of righting behavior. Thus, the pruning of inappropriate synapses is an essential constraint for the emergence of mature sensory-motor circuits and normal motor behavior.

### Complement protein C3 mediates synapse elimination in sensory-motor circuits

Our study uncovers two molecularly distinct mechanisms that are causally responsible for the elimination of inappropriate synapses. One of these involves activation of the classical complement cascade through C1q-C3 signaling. The function of C3 in synaptic pruning has also been reported to be involved in the retinogeniculate system ^49^. However, does the involvement of C3 protein results in indiscriminate pruning of synapses? The answer appears to be negative, since in the hippocampal area, C3 has been implicated in the elimination of VGluT2+ synapses, leaving VGluT1+ synapses unaffected ^50^. In our study, we demonstrated that VGluT1+ synapses are eliminated, suggesting that C3 is not responsible for the removal of selective synapses. However, it is possible that specific type of proprioceptive synapses, such as Ia, Ib or II ^12^ may be selectively targeted for elimination by C3. This observation raises the possibility that the selection of synapses to be eliminated is made by the upstream protein in the classical complement cascade, C1q. Alternatively, C3 may play a different role in synaptic elimination depending on regional differences between brain and spinal cord.

Although C3 is a downstream target of C1q, it can also be produced by the lectin and alternate pathways ^51, 52^. However, it is unlikely that the latter pathways can be involved in synaptic pruning since they are associated with infection. A previous study in the brain, demonstrated that synapse elimination is mediated by C3 activation and interaction with CD11b (C3 receptor) from microglia ^53^. In our study, C3 is involved in the elimination of inappropriate synapses similar to C1q. Intriguingly, C3-mediated synapse elimination appears to be focused on somatic synapses rather than dendritic ones. This observation may offer an insight into the way the classical complement targets synapses for elimination according to their postsynaptic site.

An important question regarding the role of inappropriate synapses relates to their function. Do inappropriate synapses transmit and if so, are they similar to the appropriate ones? Our functional assays argue that inappropriate synapses function to a similar level as the appropriate synapses. This observation indicates that C1q or C3 are not merely tagging dysfunctional or silent synapses as part of the clearance process. Instead, it suggests that inappropriate synapses might express a signal, sensed by the classical complement proteins, that is not present or detectable in the appropriate synapses.

### CD47 causes synapse elimination in the spinal cord, unlike its neuroprotective function in the brain

During the last decade, CD47 and its receptor SIRPα have been described as a “don’t eat me” signal in the central nervous system, in particular, in areas such as hippocampus, retinogeniculate circuit, and cortex ^30, 54, 55^. In striking contrast however, our study demonstrates a novel and unexpected function for CD47 within the spinal cord. Should CD47 act as a neuroprotective agent in the spinal cord, its ablation would have increased synapse elimination as reported in brain regions. Remarkably, CD47^-/-^ animals exhibited the opposite effects, with its depletion inducing the same level of synaptic pruning as genetic deletion of the classical complement proteins C3 and C1q.

A common assumption regarding the protective function of CD47 implicates SIRPα. In our study, we found that SIRPα is expressed by microglia and contacts CD47, implying a similar – if not identical - mechanism of CD47 interaction. In addition, SIRPα is present in proprioceptors, suggesting that the synapses on motor neurons destined to be pruned, might express SIRPα. However, how inappropriate synapses that contact antagonistic motor neurons are selectively pruned though CD47-SIRPα signaling remains to be elucidated. Our experiments demonstrate that CD47 is a major player in synaptic elimination of inappropriate synapses in the developing spinal cord. Although CD47 acts similarly to the classical complement proteins in eliminating inappropriate synapses, a distinct difference between the two mechanisms may be the postsynaptic site – soma vs dendrite - in which synapses are eliminated. To this end, CD47 tends to remove dendritic synapses, whereas C3 appears to be involved in the pruning of somatic synapses. The significance of our study is underlined by the unanticipated function of CD47 in synapse elimination in spinal neuronal circuits, which suggests that the classical complement cascade acts alongside with CD47, providing a steadfast fail-safe dual mechanism in the elimination of inappropriate synapses.

## Supporting information

Ext Data Fig 1 Florez-Paz Mentis

Ext Data Fig 2 Florez-Paz Mentis

Ext Data Fig 3 Florez-Paz Mentis

Ext Data Fig 4 Florez-Paz Mentis

Ext Data Fig 5 Florez-Paz Mentis

Ext Data Fig 6 Florez-Paz Mentis

Ext Data Fig 7 Florez-Paz Mentis

Video 1 Florez-Paz Mentis

Video 2 Florez-Paz Mentis

Video 3 Florez-Paz Mentis

## ACKNOWLEDGMENTS

We would like to thank Livio Pellizzoni, Joriene de Nooij and Francisco Alvarez for the critical comments on the manuscript. We are grateful for the advice of Francisco Alvarez (Emory) in the density map experiments. We also thank John Pagiazitis and Geo Ables for technical assistance and help with mouse husbandry and genotyping. GZM is supported by the following NIH grants: R01-NS078375, R01-NS125362, R01-AA027079 (NIH Blueprint for Neuroscience) and Project ALS.

## METHODS

### Mice

Mice were housed under pathogen-free conditions and all surgical procedures were performed on postnatal mice in accordance with the National Institutes of Health Guidelines on the Care and Use of Animals and approved by the Columbia Animal Care and Use Committee (IACUC). Animals of both sexes were used in this study.

### Immunohistochemistry

Dissections of the spinal cords were performed on P7 animals. Mice were anesthetized using 1.2% Avertin (300 mg/kg) by intraperitoneal injection, and transcardial perfusions were performed with PBS followed by paraformaldehyde (PFA) 4%. Post fixation with 4% PFA was performed overnight. The spinal cords were washed out with PBS and immersed in 5% agar, and transverse sections were collected from L4-L5 spinal segments using a vibratome (Leica VT1000S) at 55μm thickness. Sections were incubated in blocking serum containing 10% normal donkey serum in 0.01 M phosphate buffer saline with 0.3% Triton X-100 (PBS-T) for 1 hour. Sections were incubated overnight at room temperature, with primary antibodies at different concentrations in blocking serum. Sections were washed with PBS-T and secondary antibodies were incubated in blocking serum for 3 hours. Finally, sections were washed with PBS and mounted on glass slides using Fluoromount mounting media (Sigma F4680) and subsequently scanned at the confocal microscope.

For quadruple staining, an incubation with donkey anti-goat biotylinated antibody was performed after the incubation with primary antibodies in blocking serum for three hours. After wash with PBS-Triton, Streptavidin – 405 was added as secondary antibody following the protocol for secondary antibodies described above.

The antibody concentrations used in this study are as follows: VGluT1 polyclonal anti-guinea pig (1:2000; custom made) produced by Covance, designed against the epitope (C)GATHSTVQPPRPPPP which lies within the n-terminus of mouse VGluT1. The VGluT1 antibody was validated in VGluT1 knock-out tissue ^36^. ChAT polyclonal anti-goat (1:200; Millipore, AB144P). CD47 monoclonal anti-rat (1:500; BD Pharmingen, 555297). GFP polyclonal anti-chicken (1:500; Aves labs, 1020). Ds-red polyclonal anti-rabbit (1:200; Takara, 632496). C1q monoclonal anti-rabbit (1:1000; Abcam, AB182451). Iba1 polyclonal ant-goat (1:500; Abcam, ab5076). Iba1 polyclonal anti-rabbit (1:500. Wako, 019-19741). GFAP polyclonal anti-rabbit (1:500; Dako, Z0334). Donkey anti-goat biotylinated (1:250; Jackson Immuno Research Labs, 705065147). For secondary antibodies from Jackson Immuno Research Labs concentration used was 1:250: DyLight - 405 Streptavidin (016-470-084). DyLight – 488 donkey anti-chicken (703-545-155). Dylight – 488 donkey anti-rabbit (711-545-152). Cy3 donkey anti-rat (712-165-153). Cy3 donkey anti-rabbit (711-165-152). Cy5 donkey anti-guinea pig (7066-175-148). Cy5 donkey anti-rabbit (711-175-152). Cy5 donkey anti-chicken (703-175-155).

### Labelling of muscle selective motor neurons and proprioceptors

In order to label specifically motor neurons innervating the tibialis anterior (TA) or gastrocnemius (Gs) muscles and the proprioceptor to motor neuron synapses from antagonistic muscles, we used a modified protocol reported by Balaskas and colleagues ^37^. Briefly, two mouse strains were crossed, parvalbumin (Pv)-directed FlpO recombinase (Pv::FlpO), Jackson Laboratory (B6.Cg-*Pvalb^tm4.1(flpo)Hze^*/J) (ref. 022730), and Cre and Flp-dependent TdTomato reported mouse [Ai65 (RCFL-tdt)-D], Jackson Laboratory (B6;129S-*Gt(ROSA)26Sor*^*tm65.1(CAG-tdTomato)Hze*^/J) (ref. 021875) to obtain Pv::FlpO;Ai65 mice. These animals were crossed with C3^-/-^, C1q^-/-^, and CD47^-/-^ animals to obtain Pv::FlpO;Ai65;C1q^-/-^, Pv::FlpO;Ai65;C3^-/-^, and Pv::FlpO;Ai65;CD47^-/-^ strains respectively. Only homozygous animals for all the genes were used for experiments, and Pv::FlpO;Ai65 was used as control. Labeling of motor neurons innervating TA muscle and proprioceptors innervating Gs was performed as follows: newborn (P0-P1) pups were anesthetized with isoflurane 5% and 1liter/min of O2. 3 mm incisions were performed unilaterally in the right hindlimb at the front and back side of the hindlimb to expose the TA and Gs respectively. ∼1 μl of the virus was injected using a fine glass micropipette. The viruses injected were: in TA rAAV6/ds-CBH-GFP (Vector core of the University of North Carolina, USA) to label TA motor neurons. CAV2-Cre virus in Gs (Plateforme de Vectorologie de Montpellier, France) to flip the expression of Ai65 gene only in parvalbumin-positive Gs innervating cells; this allowed us to specifically label proprioceptors originating in a select muscle.

### Quantification of synapse number and CD47 tagging

Images from immunohistochemistry experiments were obtained using SP8 Leica confocal microscope (Leica Microsystems). Z-stacks images of 20-30 µm at 3.5 µm interval, were obtained using a 40x oil immersion objective, at 4096 x 4096 dpi resolution. Images were analyzed using ImageJ software (National Institute of Health). Quantification was performed using only neurons whose somata were completely covered by the Z-stack and only synapses on the soma were quantified. For synapses on dendrites, the first 50 μm dendritic compartment from the soma was included in the analysis. Synapses were labeled as “tagged” by CD47 only or CD47 and C1q when their protein expression colocalized with the synaptic signal, identified by VGlut1, and confirmed by an overlap using intensity profiling. Analysis of the total number of synapses on somata and dendrites and the number of synapses tagged were used to obtain the percentage of tagged synapses.

### RNAscope assay

Spinal cords were obtained at P1, P5, and P10 pups as was described for immunohistochemistry experiments. After overnight 4% post-fixation, the tissue was immersed in 10% sucrose in PBS at 4°C for 72 h. The tissue containing L4-L5 lumbar enlargement was obtained and frozen in Optimal Cutting Temperature media (OCT) and sections of 14 – 20 μm were cut in a cryostat (Leica SM3050S) and mounted on SuperFrost Plus slides (Fisher Scientific). RNAscope® Multiplex Fluorescent Detection Kit v2 (ACD BioTchne; Cat 323110) was used for detection. Slides were incubated at 60°C for 30 min and post-fixed with 4% PFA. Slides were dehydrated using ethanol dilutions of 50%, 70%, and 100% for 5 min each, followed by incubation with hydrogen peroxide (ACD BioTechne. Ref. 322335) for 10 min, after washing out with distilled H_2_O sections were embedded in 100% alcohol for 3 min. Protease Plus (ACD BioTechne. Ref. 322331) was added and slides were incubated at 40°C for 30 min. Hybridization was performed by adding a specific probe for each gene and incubating the slides for 2h at 40°C. Slides were washed with 1X wash buffer (ACD BioTechne. Ref. 310091) and incubated at room temperature overnight in 1X saline sodium citrate (SSC). The following day, slides were incubated with Hybridize AMP1, Hybridize AMP2, and Hybridize AMP3 for 30 min at 40°C each. Finally, HRP signals were developed using HRP-C1, HRP-C2, or HRP-C3 depending on the channel of each probe, and incubated for 15 min at 40°C. TSA fluorescein, TSA Cy3 or TSA Cy5 (PerkinELmer. Ref. NEL760001KT) were used to reveal the probe following company indications. To stop the hybridization, slides were incubated for 15 min at 40°C with an HRP blocker (ACD BioTechne. Ref. 323107), and mounted using Fluoromount mounting media (Sigma F4680).

For double RNAscope – immunohistochemistry assays, immunohistochemistry protocol was performed as indicated above starting with the blocking using blocking serum after the incubation with HRP blocker. Images were taken between 1 and 15 days after the experiment.

Probes used were obtained from ACD BioTchne and the protocol for dilution was followed as indicated by the company. Probes were, C1q (ACD BioTechne. Ref. 441221-C3), CD47 (ACD BioTechne. Ref. 515461-C2), and SIRPα (ACD BioTechne. Ref. 527228-C1).

### CD68 expression in microglia and volumetric analysis

We performed immunohistochemistry experiments in order to study the expression of CD68 in microglia in WT, C3^-/-^, and CD47^-/-^ mice at P5. The immunohistochemistry protocol used was the same as described above. Primary antibodies and concentrations used were: CD68 monoclonal antibody anti-rat (1:500, Bio-rad, MCA1957), Iba1 polyclonal anti-rabbit (1:500, Wako, 019-19741) ChAT, polyclonal anti-goat (1:200, Millipore, AB144P). Secondary antibodies used were: Dylight – 488 donkey anti-rat (712-475-153), Cy3 donkey anti-rabbit (711-165-152), and Cy5 donkey anti-goat (705-175-147). All 2ry antibodies were purchased from Jackson Immuno Research Labs. Images were obtained using an SP8 Leica confocal microscope (Leica Microsystems). Z-stacks images for 15-20 µm at 0.35 µm interval were obtained using a 40x oil immersion objective, at 4096 x 4096 dpi resolution. Further, the images were analyzed and processed using Neurolucida software, and 3D images were created to measure the volumetric distribution of microglia and CD68 expression inside microglia. Only microglia surrounding L4-L5 ChAT+ motor neurons were taking into consideration for analysis. A minimum of 5 microglia were analyzed per animal and CD68 expression in microglia was normalized by the total volume per cell.

### Genotype

DNA extraction was done using tail tissue incubated at 55°C for 60 min in 0.05% Proteinase K (Thermo Fisher Scientific. Ref. EO0491), diluted in tail lysis buffer containing: Tris 1M, EDTA 0.5M, NaCL 0.2M, SDS 0.07M. Samples were diluted at 1:20 in H_2_O. Genotype protocols for PV::FlpO, Ai65 (RCFL-tdt), C3^-/-^, C1q^-/-^, and CD47^-/-^ animals were followed as described in Jackson laboratory website (www.jax.org). For all animals universal PCR reaction was used as follows: 12.5 μl of GoTaq Hot Start Green Master Mix (Promega), 0.5 μl of each primer (25 μM; Sigma), and 4 μl of 1:20 diluted lysed tail DNA in a final volume of 25 µl using dH2O. For Pv::FlpO, and Ai65 (RCFL-tdt)-D and CD47^-/-^ alleles products were amplified using the following thermal cycling method: 94°C for 2 min, followed by 10 cycles of 94°C for 20 sec, 65°C for 15 sec, 68°C for 10 sec, then 28 cycles of 94°C for 15 sec, 60°C for 15 sec, 72°C for 10 sec, and followed by 72°C for 2 min. For C3^-/-^ alleles products were amplified using the following thermal cycling method: 94°C for 3 min, followed by 12 cycles of 94°C for 20 sec, 64°C for 30 sec, 72°C for 35 sec, then 25 cycles of 94°C for 20 sec, 58°C for 30 sec, 72°C for 35 sec, and followed by 72°C for 2 min. For C1q^-/-^ alleles products were amplified using the following thermal cycling method: 94°C for 4 min, followed by 35 cycles of 94°C for 1 min, 57°C for 30 sec, 72°C for 30 sec, and followed by 72°C for 10 min.

Customized primers used are listed as follows: for C1q, mutant 5’-3’ GGGGATCGGCAATAAAAAGAC, WT 5’-3’ GGGGCCTGTGATCCAGACAG, 3’-5’ common ACCAATCGCTTCTCAGGACC; with a product of WT ∼330 and mutant ∼120. For C3, mutant 3’-5’ TGGGCTCTATGGCTTCTGAG, WT 3’-5’ CACCTTACAGCACTCCCACA, 5’-3’ common GAAGTGGAAGTTGAACAAATCG; with a product of WT ∼300 and mutant ∼199. For Ai65 (RCFL-tdt)-D, in separate reactions, WT 5’-3’ ACGGGCAGTAGGGCTGAG, 3’-5’ AGCCTGCCCAGAAGACTCC, and mutant 5’-3’ GCAATAGCATCACAAATTTCA C, 3’-5’ TCTAGCTTGGGCTGCAGGT. For PV::FlpO, WT 5’-3’ GGATGCTTGCCGAAGATAAG, mutant 5’-3’ CTGAGCAGCTACATCAAC AGG, common 3’-5’ TGTTTCTCCAGCATTTCCAG. For CD47, WT 3’-5’ CACCTTACAGCACTCCCACA, mutant 3’-5’ TGGGCTCTATGGCTTCTGAG, and common 5’-3’ GAAGTGGAAGTTGAACAAATCG.

### Electrophysiology

The *ex vivo* spinal cord preparation used in this study has been described in previous publications ^35, 36^. Briefly, neonatal pups were decapitated and the spinal cord wa extracted in artificial cerebrospinal fluid (aCSF) at ∼12°C, and oxygenated with 95%O_2_/5%CO_2_. aCSF contains in mM: 128.35 NaCl, 4 KCl, 0.58 NaH_2_PO_4_.H_2_0, 21 NaHCO_3_, 30 D-Glucose, 1.5 CaCl_2_.H_2_0, and 1 MgSO_4_.7H_2_0. The spinal cord was transferred to the recording chamber with aCSF perfusion (∼10 ml/min) at room temperature (∼22°C). Suction electrodes were placed in the dorsal root or ventral root for either stimulation or recording.

The stimulus threshold was defined as the minimal current necessary to evoke a response in the ventral root in three out of five trials, with 0.2 ms of stimulation duration. The stimulus was delivered by a constant current stimulus isolator (A365 - WPI, Sarasota, FL). The recordings were amplified 1000x (Cyberamp, Molecular Devices), fed to an A/D interface (Digidata 1320A, Molecular Devices), and acquired at 50kHZ using Clampex (v10, Molecular Devices). Data were then analyzed offline using Clampfit (Molecular Devices). Only monosynaptic responses over 1mV amplitude were considered for analysis. A series of 10 stimuli were performed at 0.1, 0.2, 1 and 10 Hz. Monosynaptic amplitude was measured as the average of 10 single responses at 0.1 Hz.

The *ex vivo* spinal cord-hindlimb preparation was prepared following the same protocol for the *ex vivo* spinal cord preparation. A marked difference was the careful dissection of the hindlimb in continuity with the sciatic nerve as well as the L4 and L5 dorsal and ventral roots. The sural nerve (pure sensory) was cut and common peroneal (CP) and tibial (Tb) were intact. Concentric needle electrodes were placed within the tibialis anterior (TA) and gastrocnemius (Gs) muscles. Suction electrodes were positioned “*en passant*” on CP and Tb nerves. Ten stimuli were evoked at 0.1, 0.2 and 1 Hz frequency. The amplitude was measured from baseline to pick, and its latency was measured from the onset of the stimulus artifact to onset of the response which became higher than 3x SD of the baseline noise.

Whole-cell patch clamp recordings, were performed using *ex vivo* spinal cord-hindlimb preparation. To allow pure synaptic potentials (excitatory postsynaptic potentials, or EPSPs) to be recorded and not be contaminated by antidromic action potentials from the CP nerve (for TA motor neuron recordings) the L4 and L5 ventral roots were cut and placed in suction electrodes. The spinal cord was placed on the lateral side to access motor neurons from the lateral funiculus. The dura mater was removed between the L4-L5 area. The intracellular electrode was guided by a motorized micromanipulator, guided by fluorescence signal from motor neurons retrogradely labelled at earlier age and visualized with the aid of a Leica epifluorescent microscope (DM 6000FS). The intracellular solution contained (in mM): 130 K-gluconate, 10 NaCl, 10 HEPES, 1 EGTA, 1 MgCl_2_, 0.1 CaCl_2_, and 1 Na-GTP. pH was adjusted to 7.2 with KOH. Osmolality was adjusted to 290-295 mOsmol. Additionally, Alexa555 (1 mg/ml) and Neurobiotin (1 mg/ml) were added to the intracellular solution, as to visualize the recorded cell *post hoc*. Borosilicate electrodes were pulled using a puller P-1000 (Sutter) with resistance ranging between 13-18 MΩ. The junction potential was corrected and taken into account for subsequent analysis. The input resistance for each cell was obtained from the slope of a steady-state (linear) current-voltage plot in response to a series of hyperpolarizing currents of 50 pA each. To confirm physiologically the recorded neuron as a motor neuron, stimulation of ventral root resulted in antidromically-evoked action potentials. Signals were acquired using a Digidata 1440A (Molecular Devices., Sunnyvale, CA, USA) controller by pClamp 10.3 software. EPSC and I-V responses were analyzed off-line with the software Clampex 10.1 (Molecular Devices).

### Spatial distribution of motor neuron populations

To identify the spatial distribution of TA and Gs motor neurons, a modified protocol previously described by Bikoff and collaborators was followed ^15^. Briefly, samples were obtained from P5 animals injected with CTB488 in TA and CTB555 in Gs at P0. Slides were cut of 75 μm and immunohistochemistry against ChAT was performed in order to confirm that labeled cells were motor neurons. Confocal images were obtained from areas only between L4-L5 and images in which both TA and Gs motor neurons were located at the same space, were used for analysis. Z-stack maximal projections were obtained using Image software. Images were analyzed using Neurolucida software and seven consecutive z-stack images were used (450 μm total). Using the “LABEL” function, TA and Gs motor neurons were manually localized. Additionally, spinal cord contour was delimited, and the central canal was taken as reference point 0,0 (in the X and Y axis). Cartesian coordinates for each motor neuron were determine and obtained in reference of the central canal. Coordinates were exported as .xls files and plotted using Matlab extension “make contours”. Analysis of the density distribution was performed based on the centroid of the population. Centroid was defined as the mean x and y coordinates across cells. The mean of the cell population distribution in the ventro-dorsal and the medio-lateral aspects was obtained for each animal, and values were average for each population.

### Statistical analysis

Statistical analysis was performed using Graphpad software (PRISM version 6.04.). Comparisons were performed by either two-tailed unpaired t-test, One-Way ANOVA, using Tukey’s as *post hoc* test when conditions of normality and homoscedasticity were respected. If violated, Mann–Whitney-Wilcoxon and Kruskal-Wallis tests were used as non-parametric tests. *Post hoc* multiple comparison methods are indicated in the Results and figure legends when necessary. We presented our results in two ways: i) data points (in grey) with bars show all neurons analyzed in each group and ii) colored data points ± standard error of the mean (SEM) showing the average value measured for each mouse. Statistical comparison was performed amongst the average values from mice, in order to avoid issues of pseudoreplication in the cases that we have measured synapses from many motor neurons across different number of mice. To test for homoscedasticity (equality of variance) we performed the Brown-Forsythe test for values of individual motor neurons in each mouse. Statistical significance is provided (actual p value) in each figure legend and assigned in the graphs on the following criterial: * p ≤ 0.05, ** p ≤ 0.01, and *** p ≤ 0.001. Results are expressed as means ± standard error of the mean (SEM).

## LEGENDS FOR SUPPLEMENTARY FIGURES

**Extended Data** Figure 1**. Effects of genetic deletion of C3, CD47 or combined in the sensory-motor circuit. A)** Single optical plane confocal images of L4/5 spinal cord ventral horns from WT, C3^-/-^, CD47^-/-^ and C3^-/-^CD47^-/-^ mice at P1 (left column) and WT, C3^-/-^, C3^+/-^, CD47^+/-^, CD47^-/-^ and C3^+/-^CD47^+/-^ at P10 (middle and right columns), with antibodies against ChAT (blue), VGlut1 (green). Note that double homozygous knock out mice for C3 and CD47 (C3^-/-^CD47^-/-^) are not viable after ∼P3. Scale bar: 50 μm. **B)** Number of VGluT1 synapses per motor neuron soma in WT and C3^+/-^ (het) mice, at P10 (N=3 mice/group). Number of motor neuron somata analyzed, WT: n=17 MNs; C3^+/-^: n=16 (grey data points), from N=3 mice/group (colored data points). * p<0.05, unpaired t-test. **C)** Number of VGluT1 synapses per motor neuron soma in WT and CD47^+/-^ (het) mice, at P10 (N=3 mice/group). Number of motor neuron somata analyzed, WT: n=17 MNs; CD47^+/-^: n=13 MNs (grey data points), from N=3 mice/group (colored data points). * p<0.05, unpaired t-test. **D)** Number of VGluT1 synapses per motor neuron soma in WT, C3^+/-^, CD47^+/-^ (hets) and C3^+/-^CD47^+/-^ (double het) mice, at P10 (N=3 mice/group). Number of motor neuron somata analyzed, WT: n=17 MNs; C3^+/-^: n=16 MNs; CD47^+/-^: n=13 MNs; C3^+/-^CD47^+/-^: n=22 MNs (grey data points), from N=3 mice/group (colored data points). Significance: *p<0.05, **p < 0.001, ***p < 0.001; One-way ANOVA, Tukey’s *post hoc* test. **E)** Normalized (with respect to 1st response) amplitude change of spinal reflex challenged at 0.1Hz and 10Hz in WT (blue: 0.1Hz, N=4 mice; 10Hz, N=3), C3^-/-^ (green, 0.1Hz, N=4; 10Hz, N=4), and CD47^-/-^ mice (purple: 0.1Hz, N=4; 10Hz, N=4) at P5. Significance: C3^-/-^ vs WT: *p < 0.05; two-way ANOVA multiple comparisons with Bonferroni’s test. **I)** Righting time for WT (N=16 mice), C3^+/-^ (N=4), CD47^+/-^ (N=7), and C3^+/-^CD47^+/-^ (N=6) animals from P1 until P10. **F)** Same analysis as in (E) but at P10. WT (blue, 0.1Hz, N=8; 10Hz, N=8), C3^-/-^ (green: 0.1Hz, N=4; 10Hz, N=4), and CD47^-/-^ (purple, 0.1Hz, N=3; 10Hz, N=3). No significant difference was observed. two-way ANOVA multiple comparisons with Bonferroni’s test. **J)** Body weight gain in WT (N=16), CD47^+/-^ (N=9), C3^+/-^ (N=4) and C3^+/-^CD47^+/-^ (N=6) mice. No significance was observed, One-way ANOVA multiple comparisons with Bonferroni’s test. **H)** Body weight gain in WT (N=16), CD47^-/-^ (N=17), C3^-/-^ (N=16) and C3^+/-^CD47^+/-^ (N=6) mice. Significance: C3^-/-^ vs WT: * p<0.05, ** p<0.01, *** p<0.001; C3^+/-^CD47^+/-^ vs WT: # p<0.05, One-way ANOVA multiple comparisons with Bonferroni’s test.

**<u>Extended Data</u>** Figure 2**. Exclusion criterion for inappropriate synapses which are formed between TA proprioceptors onto Gs motor neurons and density maps for TA and Gs motor neuron pools. A)** Confocal images showing DsRed (red), ChAT (blue), VGluT1 (grey) and rAAV6/ds-CBH-GFP (green) immunoreactivity depicting a TA motor neuron (labelled by AAV6-GFP from the TA muscle) innervated by homonymous proprioceptors (induced by CAV2-Cre in TA muscle) in a PV::FlpO;Ai65 mice at P5. The magnified insert shown in the merged imaged corresponds to dotted box. Arrows point to TA proprioceptive synapses colocalized with VGluT1. **B)** Criterion for exclusion of synapses from our analysis. Image depicting an excluded synapse labelled in DsRed (red) and colocalized with VGluT1 (green) because it was not apposed to a dendrite of a TA motor neuron (shown in green, see graph where there is no overlap between red+blue with green) in a P5 PV::FlpO;Ai65 mouse at P5. **C)** Z-stack confocal images (total: 3.5 μm) from a Gs motor neuron pool in PV::FlpO;Ai65 and PV::FlpO;Ai65;C3^-/-^ mouse injected with rAAV6-GFP virus in Gs muscle and CAV2^CRE^ in the TA muscle. **D)** Percentage of motor neurons with inappropriate synapses from TA proprioceptors to Gs motor neurons. Significance: *p < 0.05; Unpaired t-test. **E)** Identification of inappropriate synapses in PV::FlpO;Ai65;C1q^-/-^mice, labelled by DsRed (red) and VGluT1 (blue) on TA motor neurons (GFP; green). Yellow arrows show dendritic and somatic inappropriate synapses in higher magnification images. **F)** Percentage of motor neurons with inappropriate synapses in C1q^-/-^ mice (WT, N=3 mice; C1q^-/-^, N=4). ** p < 0.001; Unpaired t-test. **G)** Density maps for TA (red) and Gs (blue) motor neuron pools. Maps represent the ventro-dorsal and medio-latearal distribution of MNs between L4 and L5 (450μm total). 0 (y-axis) represent central canal. **H-I)** Quantification of “hot area” distribution of each type of motor neuron, TA (H) and Gs (I). ns: no significance, One-way ANOVA multiple comparisons with Bonferroni’s test. D: dorsal, V: ventral, M: medial, L: lateral.

**<u>Extended Data</u>** Figure 3**. H-responses evoked by antagonistic nerve stimulation in C1q^-/-^ mice. A)** EMG responses from the TA and Gs muscles in response to CP nerve stimulation in a C1q^-/-^ mouse. Red traces depict the average of five responses (grey) acquired at 0.1Hz. Black arrowheads indicates stimulation artifact. Green arrow indicates the H-reflex. Blue arrow indicates the inappropriate H-response in the antagonistic muscle (magnified in the inset). **B)** Amplitude measurements of H-response following CP nerve stimulation in the TA muscle (CP nerve → TA EMG) and in the antagonistic Gs muscle (CP nerve → Gs muscle) in WT and C1q^-/-^ mice. WT: N=6 mice, C1q^-/-^: N=6. Significance: *p < 0.05; Unpaired t-test; ns: no significance. **C)** Latency of responses following CP nerve stimulation for M responses (empty circles), for H-reflex in the TA muscle (red squares) and H-response in the antagonistic Gs muscle (red circles). In one mouse, no response was evident in the Gs muscle. ns: no significance, Unpaired t-test. **D)** Superimposed EMG recordings from TA (top) and Gs (bottom) following CP nerve stimulation, revealing F-wave responses at supramaximal stimulation intensities. Five trials with the average (in red) responses are shown. At very high stimulation intensities to reveal F-waves, the strength of the muscle contraction during the M-response in the TA EMG, resulted in a movement artifact recorded in the Gs EMG, as shown by their identical latencies (a’ and a’’; dotted blue lines). In contrast, the H-response in the TA EMG is produced earlier than the inappropriate H-response in the Gs EMG (see latency a’’’). F-wave responses are evident at longer and variable latencies, compared to time locked latency of the H-response in either of the two muscles.

**<u>Extended Data</u>** Figure 4**. Intrinsic properties of TA motor neurons, CMAP latency in C3^-/-^ and CD47^-/-^ mice and morphological identification of TA motor neurons. A)** Superimposed antidromic action potentials in TA motor neuron following stimulation of the L4 ventral root. Red arrowhead indicates stimulation artifact. **B)** Resting membrane potential of recorded TA motor neurons in WT, C3^-/-^ and CD47^-/-^ mice. One motor neuron was recorded from each mouse. WT: N=7; C3^-/-^: N=5; CD47^-/-^: N=7. **C)** CMAPs’ latency from the experiments shown in Figure 4A. **D)** Firing frequency-current relation in TA motor neurons in WT, C3^-/-^ and CD47^-/-^ mice. **E)** Single optical plane confocal images of visualized filled motor neurons confirming their identity as of that innervating the TA muscle. Neurobiotin (blue) and CTb488 (green) is colocalized. Scale bar: 20 μm.

**<u>Extended Data</u>** Figure 5**. Jitter test of latency confirms monosynapticity of EPSCs. A)** Superimposed EPSC evoked at 0.1, 0.2 and 1Hz. Vertical dotted line with arrows indicate time-locked onset of EPSC at different frequencies for either homonymous (CP nerve) or antagonistic (Tibial nerve) stimulation. Recordings from a TA motor neuron of a CD47^-/-^ mouse at P5. Black arrowheads indicate stimulus artifact. Coefficient of variation for the latency of the EPSC onset in TA motor neurons following: **(B)** Common Peroneal (homonymous) nerve stimulation and **(C)** Tibial (antagonistic) nerve stimulation. ns: no significance, One-way ANOVA, Tukey’s *post hoc* test.

**<u>Extended Data</u>** Figure 6**. Effects of C1q genetic deletion on the sensory-motor circuit. A)** Single plane confocal image of VGluT1 (red), CD47 (green) and ChAT (blue) immunoreactivity in a C1q^-/-^ mouse at P5. Scale bar: 50 μm. **B)** Percentage of somatic synapses tagged by CD47 at P5 and P10 (N=3 mice/group; colored points in the graph). # synapses analyzed, WT: n=14 at P5, n=16 at P10; C1q^-/-^: n=17 at P5, n=13 at P10 (grey points) for N=3 mice/group (colored points represent average values per mouse). **C)** Percentage of dendritic synapses tagged by CD47. Number of dendrites; WT: n=24 at P5, n=30 at P10; C1q^-/-^: n=24 at P5, n=30 at P10 (grey points) for N=3 mice/group (colored points represent average values per mouse). Significance: * p<0.05; One-way ANOVA multiple comparisons with Bonferroni’s test. **D)** Confocal images of L4/5 spinal cord ventral horns from C1q^-/-^ mice at P1, P5 and P10 with antibodies against ChAT (blue) and VGlut1 (white). Scale bar: 50 μm. **E)** Number of VGluT1 synapses per motor neuron soma at P1, P5 and P10 mice (N=3 mice/group; colored points in the graph) in WT and C1q^-/-^ mice. Number of motor neuron somata, WT: n=18 at P1, n=14 at P5, n=16 at P10; C1q^-/-^: n=12 at P1, n=17 at P5, n=13 at P10. *p<0.05, **p<0.01, One-way ANOVA, Tukey’s *post hoc* test. **F)** Synaptic density (# synapses/50μm dendrite) for proximal dendrites of motor neurons (N=3 mice/group; colored points in the graph). Number of dendrites, WT: n=30 at P1, n=24 at P5, n=30 at P10; C1q^-/-^: n=30 at P1, n=24 at P5, n=30 at P10. ns: no significance. **G)** Electrical responses from ventral roots following stimulation of the L4 (or L5) dorsal root in WT and C1q^-/-^ mice. Red traces shown the average of first 5 responses (grey) acquired at 0.1Hz. Arrowhead indicates stimulus artifact. The dotted line indicates baseline. Blue arrows indicate peak amplitude of the averaged response. **H)** Amplitude and latency of spinal reflexes at P5 in WT (blue, N=6 mice) and C1q^-/-^ (red, N=5) preparations. Significance: *p < 0.05, Unpaired t-test.

**<u>Extended Data</u>** Figure 7**. No changes in CD47 expression in motor neurons in C3^-/-^ mice and interaction between CD47 and SIRPα on proprioceptive synapses. A)** Confocal images showing CD47-mRNA (RNAscope, in green) and immunoreactivity signals with ChAT (blue), Iba1 (red, top row), GFAP (red, bottom row) and DAPI (white) in C3^-/-^ mice at P5. Dotted boxes indicate areas of higher magnification shown on the right. Scale bar: 20 μm. **B)** Quantification of CD47 puncta per soma in WT (P5, N=4; P10, N=5) and C3^-/-^ (P5, N=5; P10, N=5). ns = no significance; One-way ANOVA multiple comparisons with Bonferroni’s test. **C)** Single plane confocal images of Parvalbumin (green), CD47 (red), and SIRPα (blue) in a WT mouse at P5. Merged image shows interaction of CD47 and SIRPα on proprioceptive (Pv+) dendrites. White arrows point to CD47–SIRPα interaction. **D)** Confocal images at low magnification of the ventral horn showing ChAT (green), CD47 (red), and SIRPα (blue) immunoreactivity. **E_1_)** Higher magnification image (dotted box in D), showing interaction (white arrows) between CD47 and SIRPα on dendrites **(E_2_)** and soma **(E_3_)** of motor neurons.

