## Supplementary figures and images for "C3 and CD47 mediate sensory-motor circuit refinement during spinal cord development"

### Ext Data Fig 1 Florez-Paz Mentis

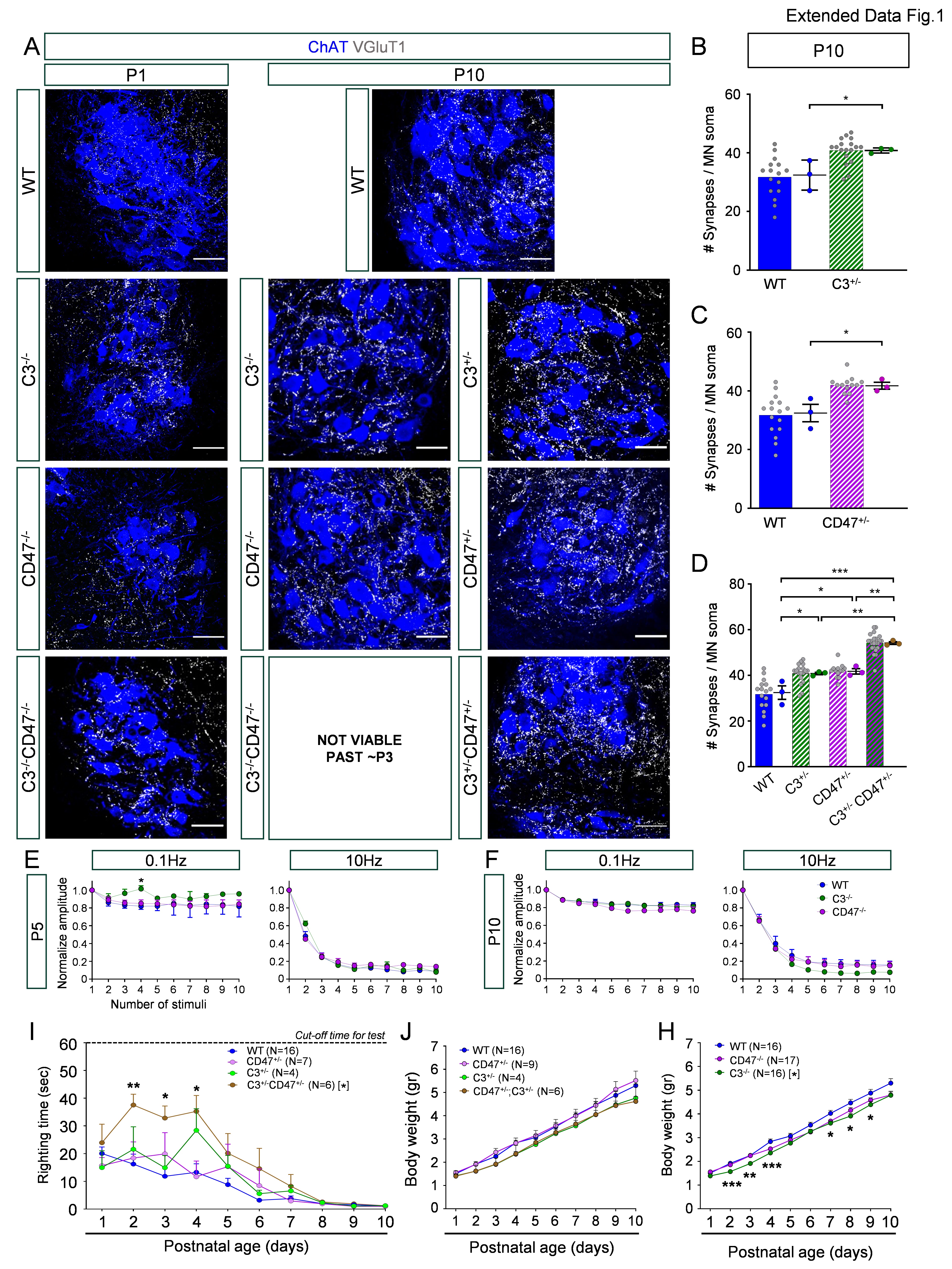

### Ext Data Fig 2 Florez-Paz Mentis

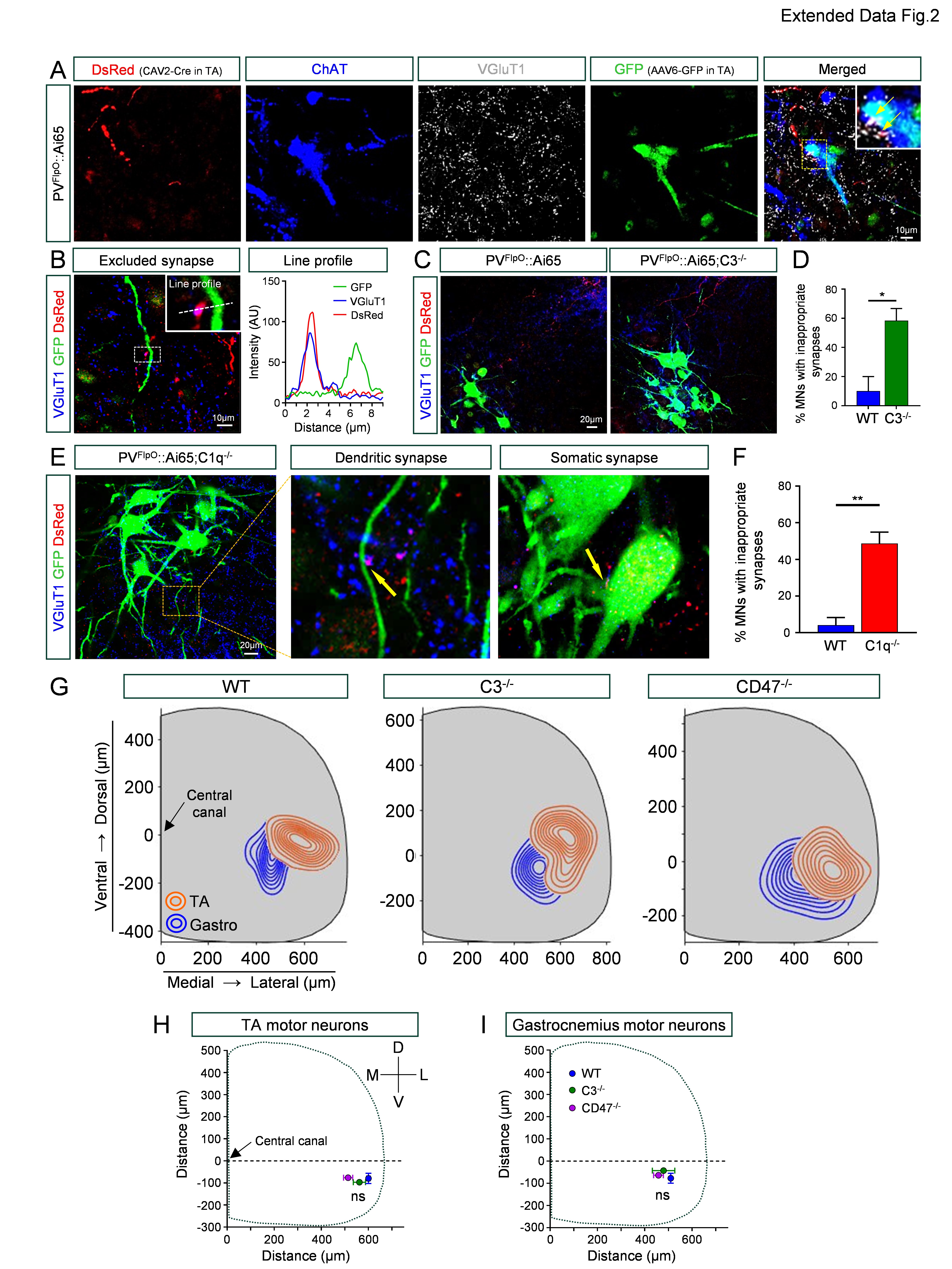

### Ext Data Fig 3 Florez-Paz Mentis

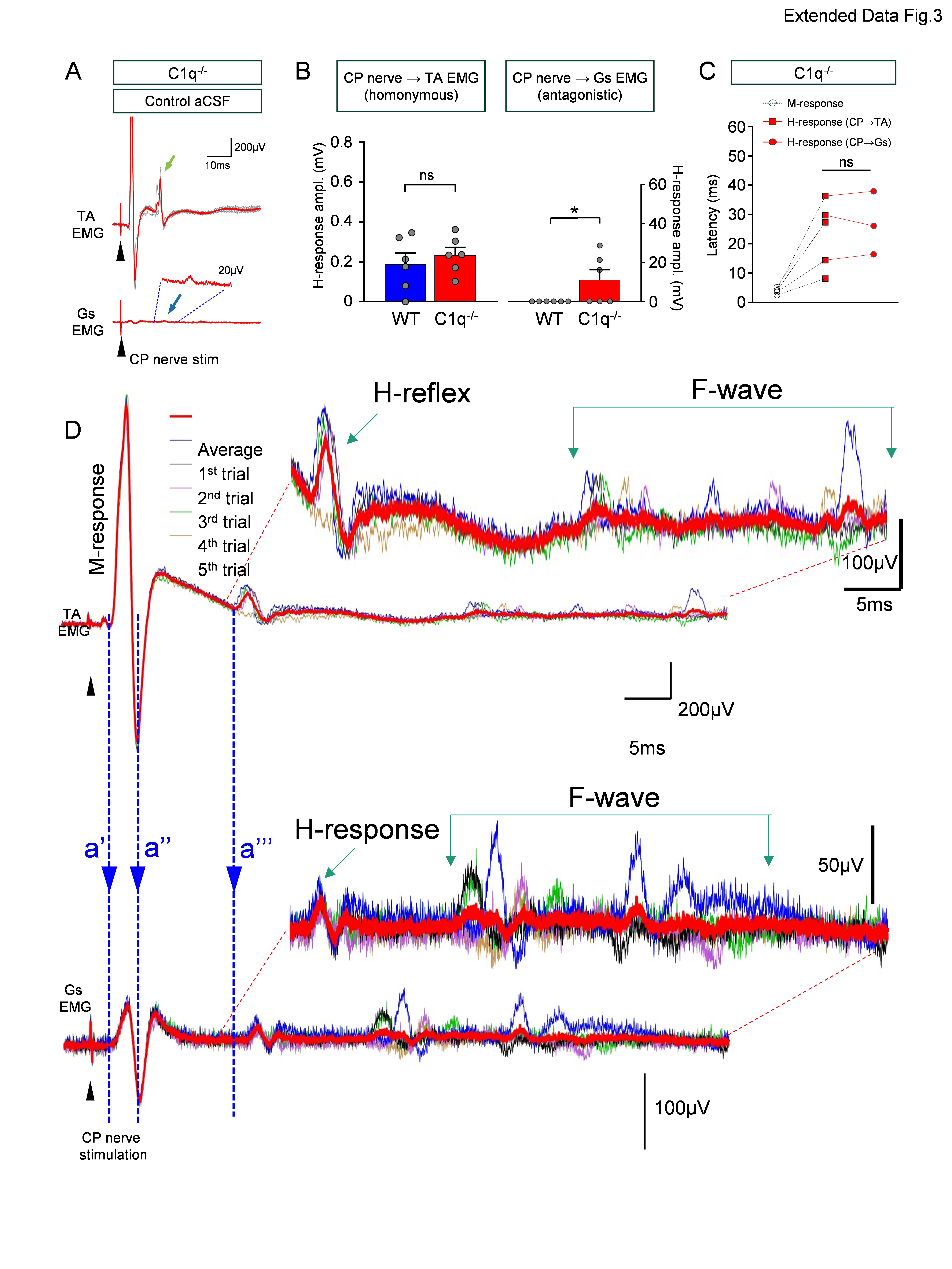

### Ext Data Fig 4 Florez-Paz Mentis

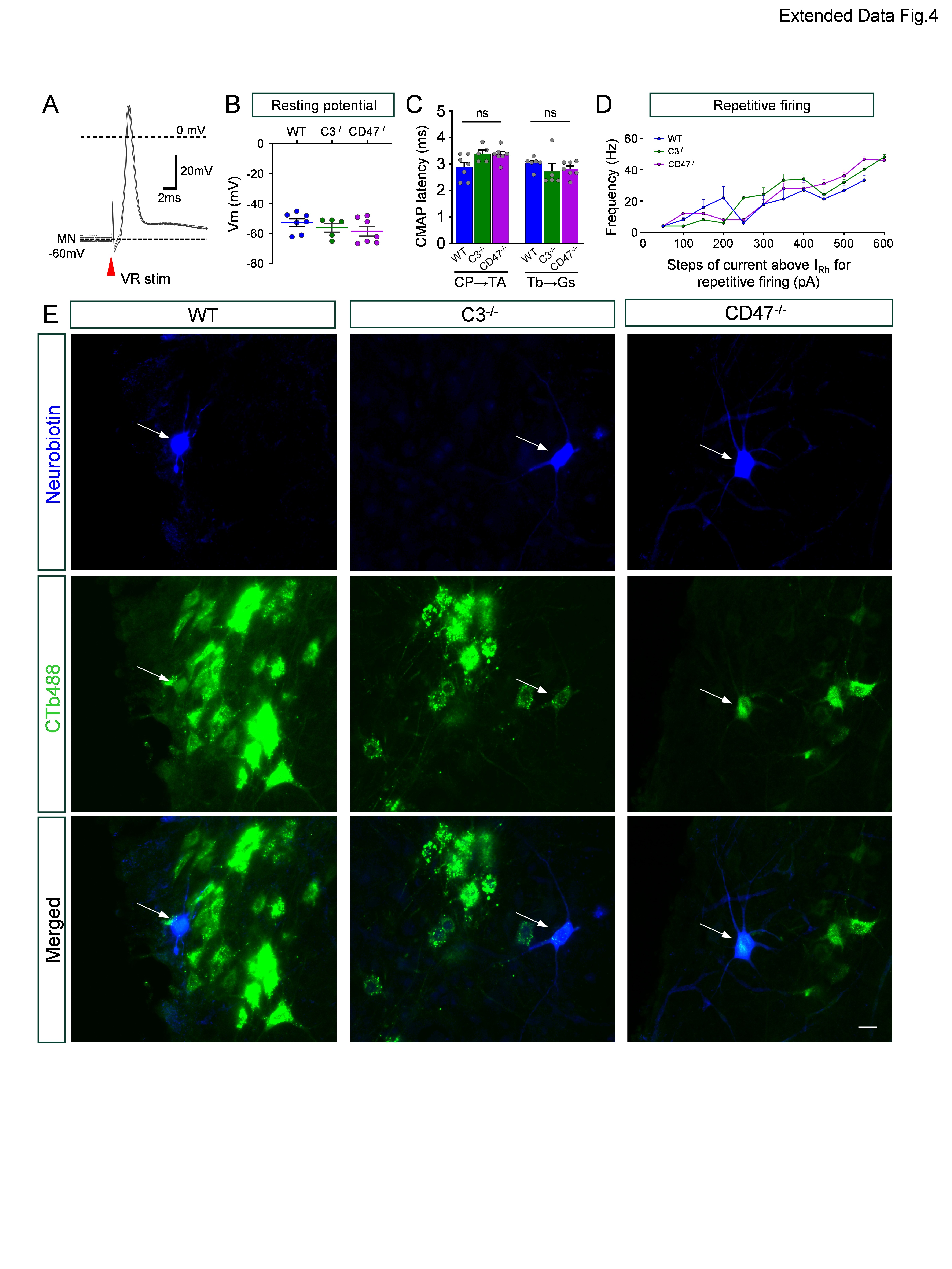

### Ext Data Fig 5 Florez-Paz Mentis

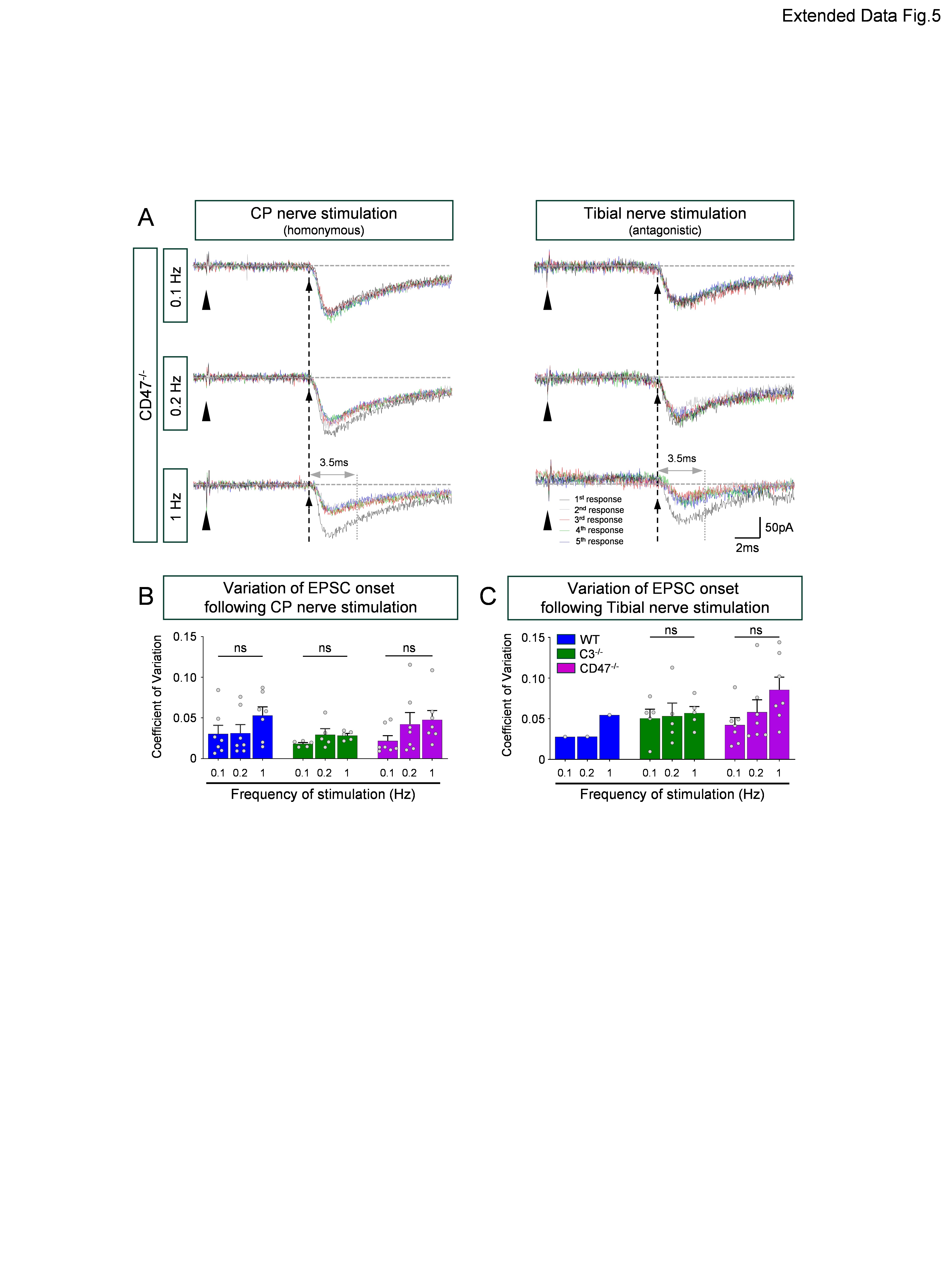

### Ext Data Fig 6 Florez-Paz Mentis

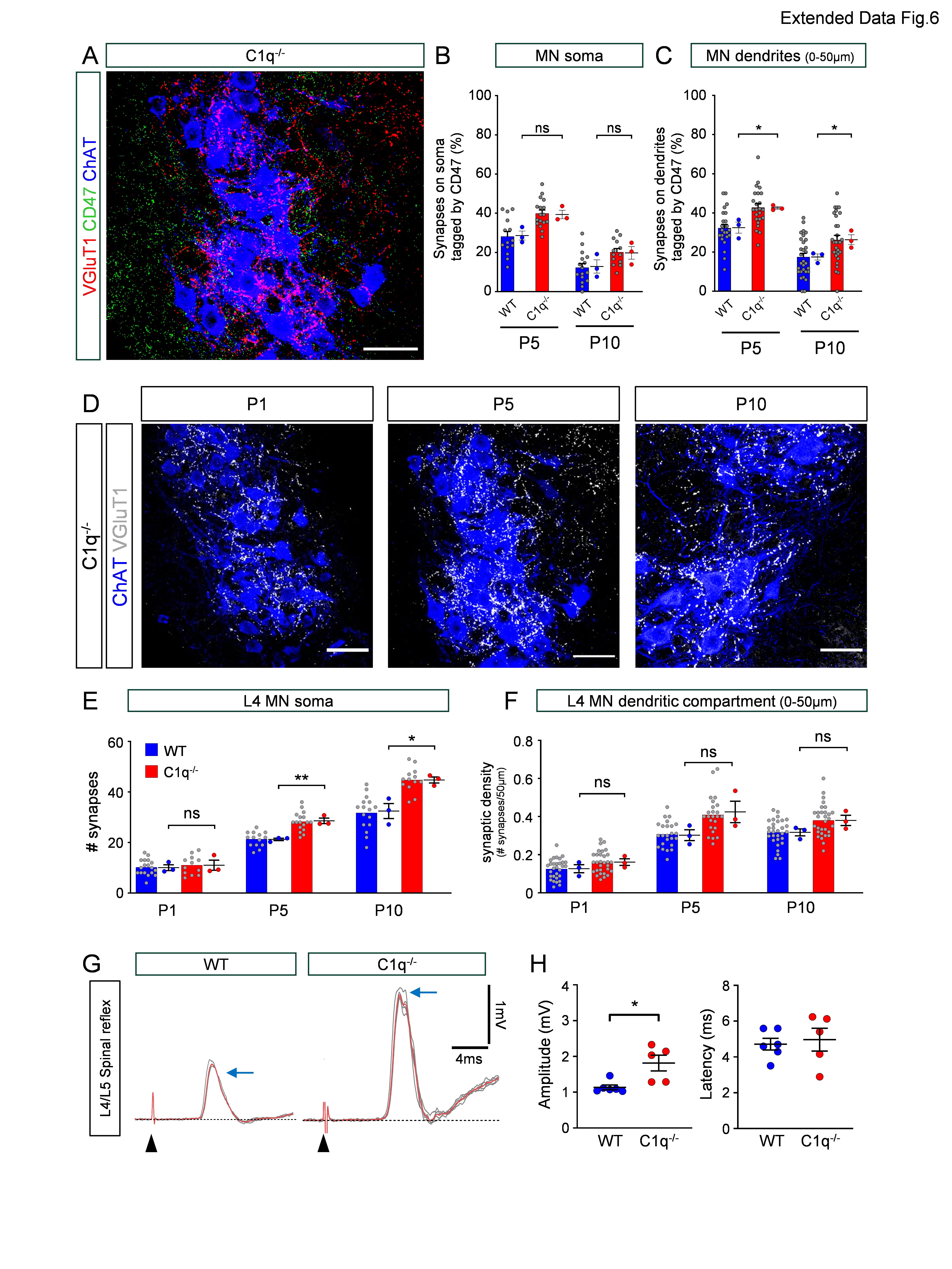

### Ext Data Fig 7 Florez-Paz Mentis

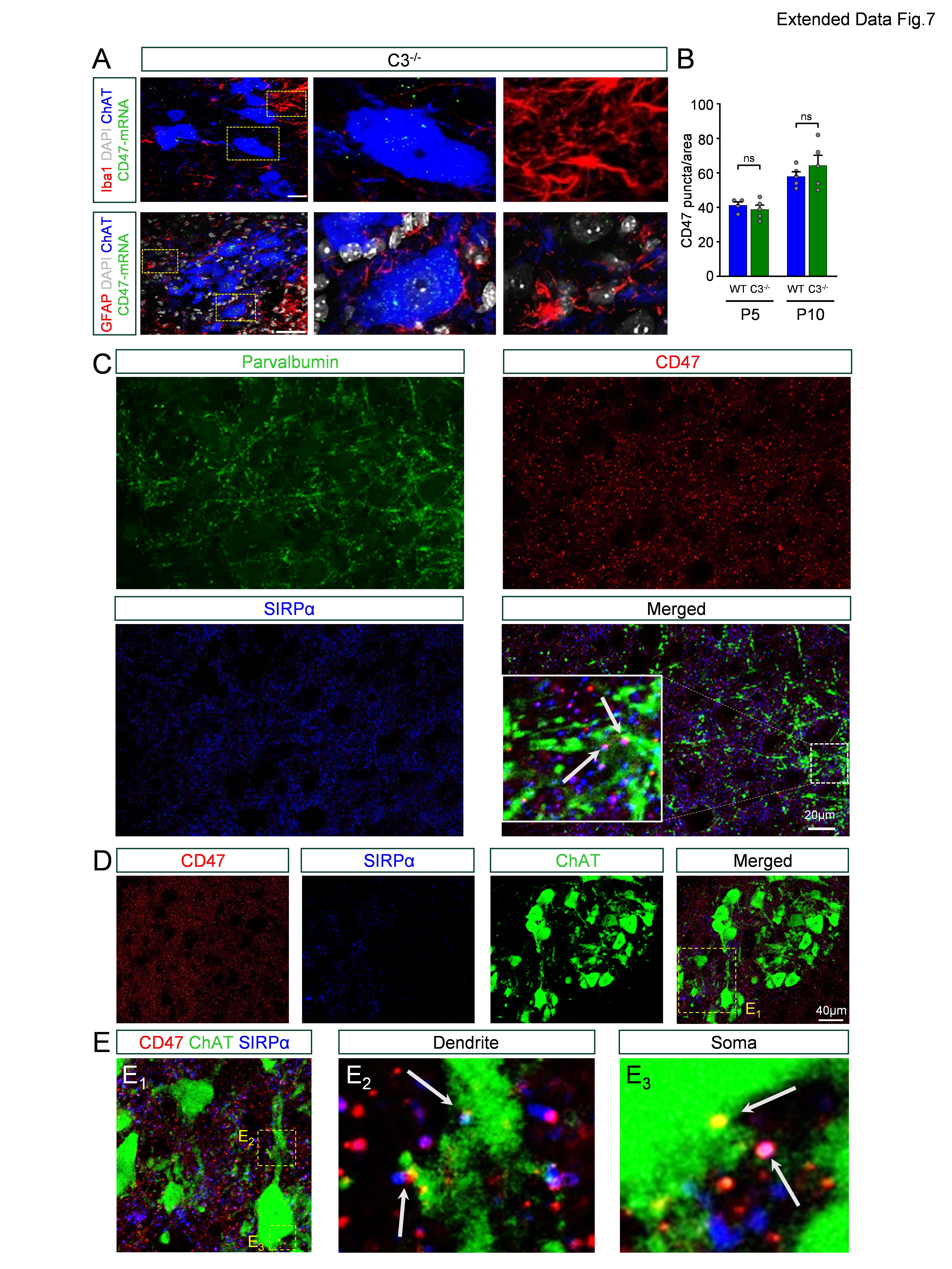
